# Phylogenomic analysis of Uganda influenza type-A viruses to assess their relatedness to the vaccine strains and other Africa viruses: a molecular epidemiology study

**DOI:** 10.1101/2021.07.05.451078

**Authors:** Grace Nabakooza, David Collins Owuor, Zaydah R. de Laurent, Nicholas Owor, John Timothy Kayiwa, Daudi Jjingo, Charles Nyaigoti Agoti, David James Nokes, David Patrick Kateete, John Mulindwa Kitayimbwa, Simon David William Frost, Julius Julian Lutwama

## Abstract

**Background:** Genetic characterisation of circulating influenza viruses is essential for vaccine selection and mitigation of viral transmission. The current scantiness of viral genomic data and underutilisation of advanced molecular analysis methods on influenza viruses circulating in Africa has limited their extensive study and representation in the global influenza ecology. We aimed to sequence influenza type-A viruses (IAVs) that previously circulated in Uganda and characterised their genetic relatedness to the vaccine viruses and publicly available Africa IAVs.

**Methods:** This was an observational study nested to the Uganda national influenza surveillance programme. We used Next-generation sequencing to locally generate genomes from 116 A(H1N1)pdm09 and 118 A(H3N2) viruses collected between 2010 and 2018 from 7 districts across Uganda. A total of 206 hemagglutinin (HA), 207 neuraminidase (NA), and 213 matrix protein (MP) sequences were genetically compared to the WHO-recommended vaccines and other viruses isolated from Africa since 1994. Viral temporal and spatial divergence and circulating genetic clades were characterised using phylogenetic methods.

**Findings:** We successfully generated gene sequences for 91·9% (215/234) viruses. Uganda A(H1N1)pdm09 and A(H3N2) virus HA, NA, and MP proteins had 96·36-99·09%, 96·49-99·39%, and 97·48-99·95% amino acid similarity, respectively, to vaccines recommended from 2010 through 2020. The local viruses incorporated amino acid substitutions (AAS) in their antigenic, receptor binding, and glycosylation sites each year causing them to antigenically drift away from vaccines. For seasons when vaccine formulations differed, Uganda IAV antigenic sites had 1-2 extra AAS relative to the Southern than Northern hemisphere vaccine viruses. All Uganda IAVs carried the adamantine-resistance marker S31N but not the neuraminidase inhibitor (NAI) resistance markers H274Y and H275Y. However, some A(H1N1)pdm09 viruses had permissive substitutions V234I, N369K, and V241I typical of NAI-resistant viruses. The 2017-2018 A(H1N1)pdm09 viruses belonged to global genetic clade 6B.1, while the A(H3N2) viruses isolated in 2017 belonged to clades 3C.2a and 3C.3a. Uganda IAVs obtained before 2016 clustered distinctly from other Africa viruses while later viruses mixed with other Africa, especially Kenya and Congo, and global viruses. Several unique viral lineages (bootstrap >90) persisted in Uganda and other countries for 1-3 years.

**Interpretation:** The study reveals Uganda as part of the global influenza ecology with continuous importation, antigenic drift, and extensive local transmission of IAVs, presenting a potential risk of future outbreaks. For a country with limited health resources and where social distancing is not sustainable, viral prevention by vaccination should be prioritized. The notable viral diversity in Africa is a warning to countries to broaden and incorporate genome analysis in routine surveillance to monitor circulating and detect new viruses. This knowledge can inform virus selection for vaccine production and assist in developing cost-effective virus control strategies.

**Funding:** This work was supported by the Makerere University-Uganda Virus Research Institute Centre of Excellence for Infection and Immunity Research and Training (MUII). MUII is supported through the Developing Excellence in Leadership, Training and Science (DELTAS) Africa Initiative (Grant no. 107743). The DELTAS Africa Initiative is an independent funding scheme of the African Academy of Sciences (AAS), Alliance for Accelerating Excellence in Science in Africa (AESA), and supported by the New Partnership for Africa’s Development Planning and Coordinating Agency (NEPAD Agency) with funding from the Wellcome Trust (Grant no. 107743) and the UK Government. The work was also funded in part by a Wellcome Trust grant (102975).

## INTRODUCTION

Novel influenza type-A viruses (IAVs) cause human respiratory infections that lead to social lockdowns, economic losses, and millions of deaths^1^. Genetic characterisation of IAVs is important to differentiate them from other viruses causing similar clinical symptoms for effective viral control and prevention. Seasonal influenza-related illnesses kill 290000-650000 people globally per year, mostly in sub-Saharan Africa^2^. Influenza accounts for 21·7% and 10·1% of the influenza-like illnesses (ILI) and severe acute respiratory illnesses (SARI) in Africa, respectively, and circulates all-year-round with discernible influenza peaks in North and South Africa^3^. Uganda’s annual epidemics have two major peaks between May and November and usually constitute multiple IAV types and subtypes responsible for 13% and 6% of the ILI and SARI cases, respectively^4^.

Vaccination and antiviral treatment are the best ways to prevent and control viral transmission^5^. However, the multi-segmented IAVs continuously mutate, especially in the antigenic surface genes, hemagglutinin (HA) and neuraminidase (NA), causing them to drift from vaccines with possible emergence of drug-resistant viruses^6^. Vaccine formulations for the Northern (NH) and Southern hemisphere (SH) are updated annually to match circulating viruses^5^. Countries select appropriately licensed vaccines based on their viral circulation patterns^5^.

Well-sampled and developed collect sufficient data on IAV evolution patterns, drug sensitivity^6^, and circulating and emerging genetic clades^7^ for vaccine selection^5^. However, the high cost of whole-genome sequencing (WGS) and limited phylogenomic analysis capacity restricts the genetic characterisation of IAVs in Africa. We searched Global Health, PubMed, and Web of Science on 25^th^ March 2020 and found only five African studies^8–12^ had sequenced IAV whole genomes (WGs). Two of these sequenced 59 A(H3N2)^8^ and 19 A(H1N1)pdm09^9^ Uganda viruses, sampled in 2008-2009 and 2009-2011, respectively, with sequencing conducted in the United States.

We aimed to explore the feasibility of using Next-Generation Sequencing (NGS) in a low-income setting to generate WGs of Uganda IAVs sampled in 2010-2018 and compare the generated sequences with vaccine viruses and publicly available Africa IAV sequences collected in 1994-2019. We analysed HA carrying antigenic sites which trigger host immune responses and NA and matrix protein (MP) genes targeted by antiviral drugs. This work birthed an East African network of influenza molecular epidemiologists which we hope to expand across Africa.

## METHODS

### Design of influenza surveillance and source of swabs

The Uganda Virus Research Institute (UVRI-NIC) National Influenza Centre in Entebbe implements a clinic and hospital-based surveillance at thirteen peri-urban and densely populated sites in seven districts located in Northwest, Western, Central, and Eastern Uganda (sFigure 1, Appendix pp 6)^4^.

Nasal and oropharyngeal swabs were collected in 2010-2018 seasons from ILI and SARI patients of all ages with their respective sociodemographic data^4^. The swabs were tested, typed for influenza A and B, and the IAV positives subtyped for seasonal [A(H1N1) and A(H3N2)] and pandemic A(H1N1)pdm09 influenza using the Centers for Disease Control and Prevention’s (CDC) real-time reverse-transcription polymerase chain reaction (rRT-PCR) protocols and primers (Atlanta, Georgia)^13^.

### Swab selection for whole-genome sequencing (WGS)

We aimed to sequence 100 whole genomes (WGs) per subtype [A(H1N1)pdm09 and A(H3N2)] due to financial constraints. Available swabs from 971 IAV patients with real-time PCR cycle threshold (CT≤35) were randomised, stratified by subtype and year of collection, and selected every first fifteen swabs. Twenty-four percent (234/971) of the swabs [116 A(H1N1)pdm09 and 118 A(H3N2)] were sequenced at a KEMRI-Wellcome Trust Programme collaborating laboratory in Kilifi, Kenya.

### Viral RNA isolation and amplification

Viral ribonucleic acid (RNA) was extracted from 140μL swab sample using the QIAamp Viral RNA Mini extraction kit and manufacturer’s protocol (Qiagen, Hilden, Germany). The extracted RNA was reverse transcribed and whole genome amplified using the multi-segment real-time polymerase chain reaction (M-RTPCR)^14^ and universal IAV Uni/Inf primers at standardised thermocycling conditions (Appendix pp 3)^11^.

### Next-generation sequencing

The M-RTPCR amplicon libraries were prepared using the Nextera XT DNA library preparation kit and protocol (Illumina, San Diego, California, USA), cleaned using the 0.8× AMPure XP beads, quantified on the Qubit 3.0 fluorometer using the dsDNA High sensitivity kit (Invitrogen, Carlsbad, California, USA). Library size distributions were assessed using the Agilent Technology 2100 Bioanalyzer and the High Sensitivity DNA kit (Agilent Technologies, Santa Clara, California, USA). Samples with a broad fragment size spectrum (>250 bp) were normalized manually to 2nM. 5μL per sample library were pooled, denatured using Sodium Hydroxide (NaOH), and diluted to 12.5 pmol. Diluted libraries were spiked with 5% Phi-X control (Illumina, San Diego, CA, USA) and sequenced using the Illumina MiSeq (Illumina Inc., San Diego, California, USA) generating 2×250bp paired reads per sample.

We assessed sequencing efficiency based on the read (coverage depth) and gene segment (genome length) count generated per swab sample.

### Sequence quality control and assembly

Raw reads were de-duplicated and decontaminated, and clean reads assembled using the reference-based FLU module of the Iterative Refinement Meta-Assembler (IRMA) v0.6.7^15^ at default settings (Appendix pp 4-5). The A(H1N1)pdm09 and A(H3N2) virus assemblies were compared with A/California/7/2009 and A/Perth/16/2009 vaccine viruses, respectively.

### Genetic characterization of surface and matrix proteins

Gene sequences for SH and NH or both (SNH) vaccine and clade reference viruses per subtype were downloaded from the Global Initiative on Sharing All Influenza Data (GISAID) EpiFlu database (https://www.gisaid.org/; accessed on 26^th^ February 2020). Uganda and vaccine virus sequences per subtype were aligned using MUSCLE v3.8.1551^16^.

Amino acid substitutions (AAS) in the antigenic sites (A, B, C, D, and E) of A(H1N1)pdm09^17^ and A(H3N2)^18^ virus HA1 proteins were identified manually. AAS in complete HA, NA, and MP protein sequences and their respective functions were identified using the Flusurver tool (http://flusurver.bii.a-star.edu.sg; accessed on 30^th^ March 2021).

### Phylogenetic divergence

Uganda HA, NA, and MP gene sequences per subtype were aligned and maximum likelihood (ML) trees reconstructed using IQtree v1.6.11^19^ with a GTR+G4 model and 1000 bootstraps. Trees were rooted using the oldest sequence in the dataset and visualized in ggtree v2.4.1^20^ and Figtree v1.4.4 (http://tree.bio.ed.ac.uk/software/figtree/).

### Phylogenetic clustering and genetic clade classification

Uganda, clade reference, and vaccine virus sequences per subtype were aligned and ML trees reconstructed as above. Phylogenetic clusters were inferred from ML trees using PhyCLIP v2.0^21^ and clusters confirmed as genetic clades using the European Centre for Disease Prevention and Control (ECDC) reference-based method^7^. PhyCLIP uses linear integer programming to cluster sequences that may share epidemiological patterns, while the ECDC method classifies a sequence or cluster of sequences to a genetic clade based on AAS shared in their HA1 or HA2 proteins.

For Africa analysis, HA, NA, and MP gene sequences were downloaded from GISAID accessed on 27^th^ February 2020. Sequences with >100 bps shorter or longer than the actual gene size and ambiguous “N” bases were excluded. The remaining A(H1N1)pdm09 (496 H1, 443 N1, and 278 MP) and A(H3N2) (718 H3, 675 N2, and 439 MP) virus sequences were aligned using a codon-aware aligner (https://github.com/veg/hyphy-analyses/tree/master/codon-msa) and ML trees reconstructed as above.

### Statistical analysis

Patients’ sociodemographic data were compared between the UVRI-NIC and study using chi-square and Wilcoxon t-test in R v3.6.3 (https://www.r-project.org).

### Ethics

This study was approved by the Makerere University School of Biomedical Sciences Research and Ethics Committee (SBS-REC) (ref:SBS-577) and Uganda National Council of Science and Technology (UNCST) (ref:HS2519).

## RESULTS

### Sociodemographic characteristics of sampled patients

The UVRI-NIC laboratory tested 18353 patients between 22^nd^ October 2010 and 9^th^ May 2018. Thirteen-percent (2404/18353) were positive for influenza, 69·88% (1680/2404), 29·62% (712/2404), and 0·17% (4/2404) had influenza A, B, and A/B co-infection, respectively. Eight (0·33%) viruses lacked viral type data (sFigure 2, Appendix pp 7). IAV positives included 67·08% (1127/1680) A(H3N2), 32·2% (541/1680) A(H1N1)pdm09, 0·12% (2/1680) AH1/H3 co-infections, and 0·59% (10/1680) lacked subtype data.

On average, 13 swabs were sampled per subtype per year except 2012, 2016, and 2018 with 2, 1 A(H1N1)pmd09, and no A(H3N2) patient swabs avaliable, respectively (sTable 1, Appendix pp 8). Three A(H1N1)pdm09 and one A(H3N2)] sampled patients’ swabs lacked sociodemographic data. Of the 230 swabs with data, 65·22% (150/230) and 34·78% (80/230) were from ILI and SARI cases, respectively. The number of ILI and SARI cases sampled per subtype were comparable (Figure 1A and 1B). More viruses were sequenced from males 56·09% (129/230) than females 43·16% (101/230). IAVs from under 5-year-olds, 72% (166/230), were most sequenced and 0·87% (2/230) from the elderly (≥50 years old) (sTable 2, Appendix pp 8). The ILI, SARI, male, and female counts were independent (chi-square test p>0·05) and patients’ mean ages not significantly different (Wilcoxon test p>0·05) between the sampled and UVRI-NIC per subtype (sFigure 3 and 4, Appendix pp 9).

**Figure 1:**
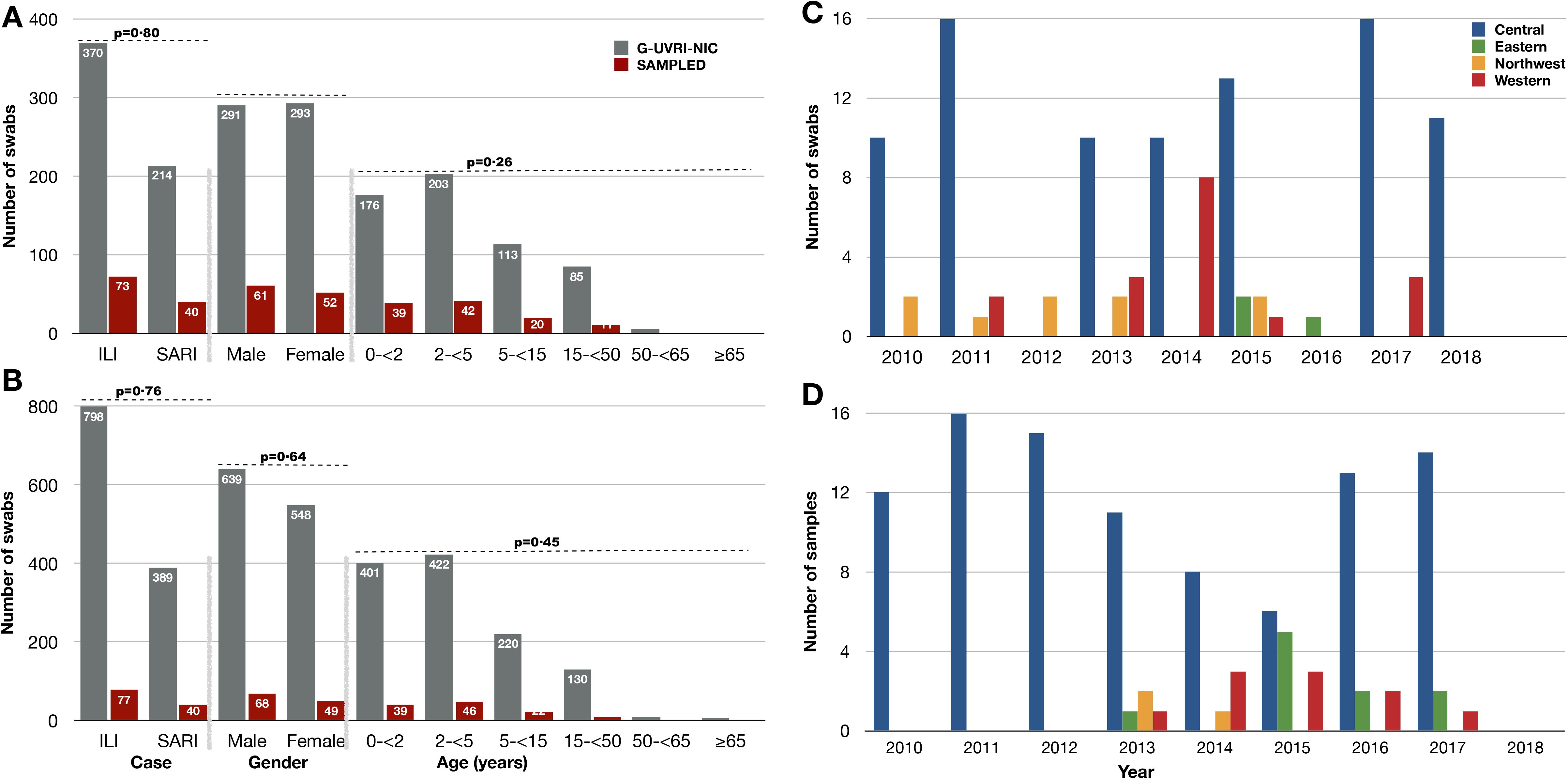
Sociodemographic characteristics of patients whose swabs were sampled for whole-genome sequencing (n=230, isolated in 2010-2018) compared to the patients sampled in the general UVRI-NIC surveillance programme in 2007-2018 (N=1800). Cases include influenza-like illnesses (ILI) and severe acute respiratory illnesses (SARI). Panel **A** shows the number of cases, gender, and different age groups (years) infected with A(H1N1)pdm09 viruses. Panel **B** shows the number of cases, gender, and different age groups (years) infected with A(H3N2) viruses. The reported p-values (chi-square and Wilcoxon) were obtained using combined cases, gender, and ages (with no age groups) per subtype. Our sampling depicts similar characteristic trends as in the general UVRI-NIC surveillance EpiInfo data. Two of the 1802 influenza A positives patients in the general UVRI-NIC surveillance dataset were co-infections of AH1/AH3, therefore only the remaining 1800 patients are included in these comparisons. **Panels C** and **D** show, respectively, the number of patients infected with A(H1N1)pdm09 and A(H3N2) viruses whose swabs were sampled and sequenced from each geographical region per year. As of 9th May 2018, the laboratory had no positive case of A(H3N2) virus.

The majority, 77·35% (181/234) of swabs sampled for WGS were from Central (Kampala and Entebbe), 11·54% (27/234) from Western (Mbarara and Fort Portal), 5·56% (13/234) Eastern (Tororo), and 5·13% (12/234) Northwest (Arua and Koboko) [Figure 1C and 1D) as in UVRI-NIC sTable 3, Appendix pp 10)].

### Sequencing efficiency

All sampled 234 swabs were analysed and read counts reported as averages.

The MiSeq generated 569435 paired reads per sample (data not shown). Following quality control (QC), 266020 (70229-909340) clean reads per sample were put in IRMA, 265868 (70150-908946) passed IRMA’s QC, 213809 (1381-908234) matched flu references, and 113164 (777-461172) paired reads were used to build assemblies (Figure 2A).

**Figure 2:**
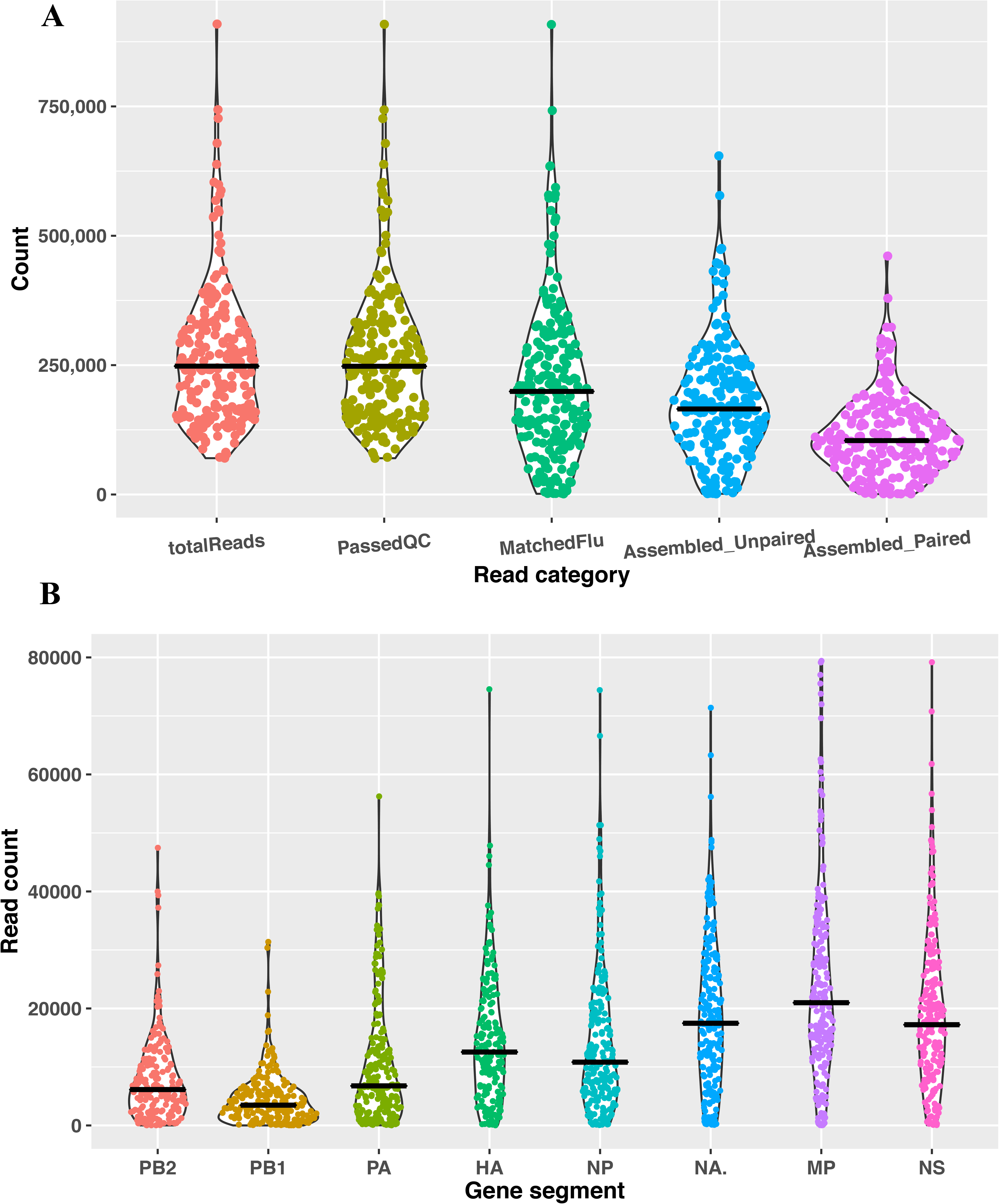
Metrics for clean MiSeq reads used for sequence assembly using IRMA Flu module. Panel **A** shows the number of reads in each read category obtained per viral swab sample. **totalReads** is the number of reads used as input for IRMA. **PassedQC** are the reads that passed IRMA internal quality control (QC). **MatchedFlu** are reads that matched IRMA’s flu references. **Assembled_Paired** are the number of paired reads used for the final assemblies, and **Assembled_Unpaired** are the number of unpaired reads used for the final assemblies. Panel **B** shows the number of paired reads used in the final assembly for each of the eight gene segments: polymerase subunits (PB2, PB1 and PA), hemagglutinin (HA), nucleoprotein (NP), neuraminidase (NA), matrix protein (MP) and non-structural protein (NS). Read metrics are for the 215 A(H1N1)pdm09 and A(H3N2) viruses retained after excluding segments with low transverse coverage depth (<100).

The number of assembled reads decreased with an increase in gene segment size. The shortest, MP and non-structural protein (NS), had 25101 (28-91585) and 19341 (41-79167) assembled reads, respectively. The NA, HA, and nucleoprotein (NP) had 18653 (74-71377), 14466 (35-74550), and 14151 (69-74415) reads assembled, respectively. The polymerase subunits: PA, PB2, and PB1 had 10465 (31-56243), 7885 (15-47444), and 4497 (8-31354) reads assembled, respectively (Figure 2B).

We successfully sequenced and assembled viruses from 96·58% (226/234) of the swabs (Figure 3). Eleven WGs (88 segments) with low coverage depth (<100) were excluded remaining with 215 viruses. 89·77% (193/215) of these were WGs, spanning 100% and >96·7% nucleotides in the coding sequences (CDS) and complete genome of A/California/7/2009(H1N1pdm09) and A/Perth/16/2009(H3N2) vaccine viruses. The remaining 10·23% (22/215) viruses had partial genomes with complete CDS for 2-7 segments. The overall WG recovery rate was 85·4% (193/226). Two viruses sampled as A(H1N1)pdm09 matched IRMA’s A(H3N2) references and were included in the A(H3N2) analysis.

**Figure 3:**
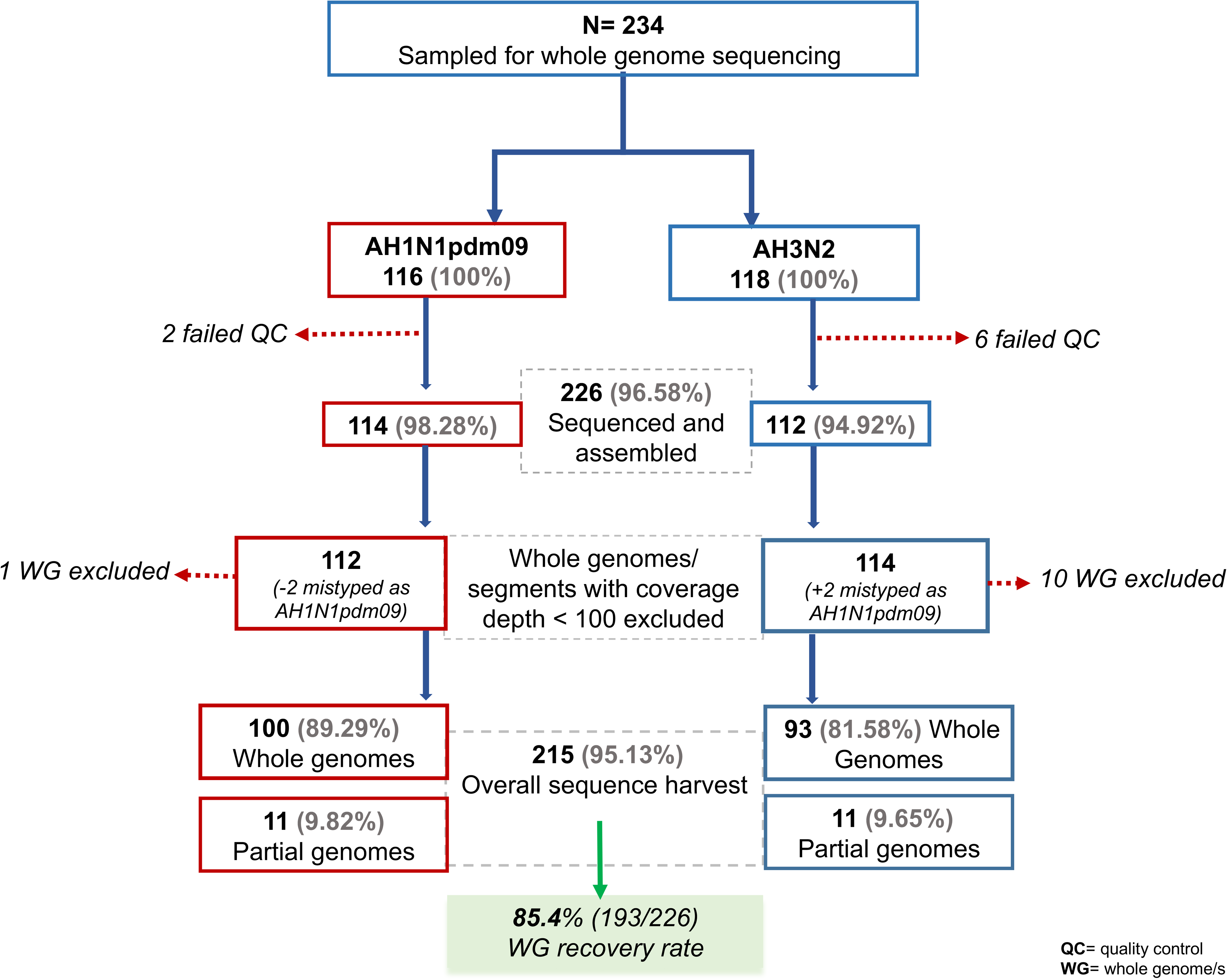
Workflow showing how viral samples were excluded before and after sequencing and the rate of whole genome recovery. Eight viruses [2 for subtype A(H1N1)pdm09 and 6 for subtype A(H3N2)] failed quality control (QC) before the sequencing.

The newly-generated sequences were deposited in GISAID under the accessions EPIISL498819-EPIISL498931 [A(H1N1)pdm09] and EPIISL498934-EPIISL499037 [A(H3N2)].

### Antigenic drift in Uganda viruses

Uganda IAV HA1 proteins continuously drifted away from 2010-2020 season vaccines (Table 1). In 2012, 2013, 2019, and 2020 when formulations differed, Uganda viruses carried 1-2 extra AAS relative to SH than NH vaccines viruses. Since Uganda’s largest part lies north of the equator, the substitutions described below are relative to NH and SNH vaccines for the sampled 2010-2018 [A(H1N1)pdm09] and 2010-2017 [A(H3N2)] seasons.

**Table 1:**
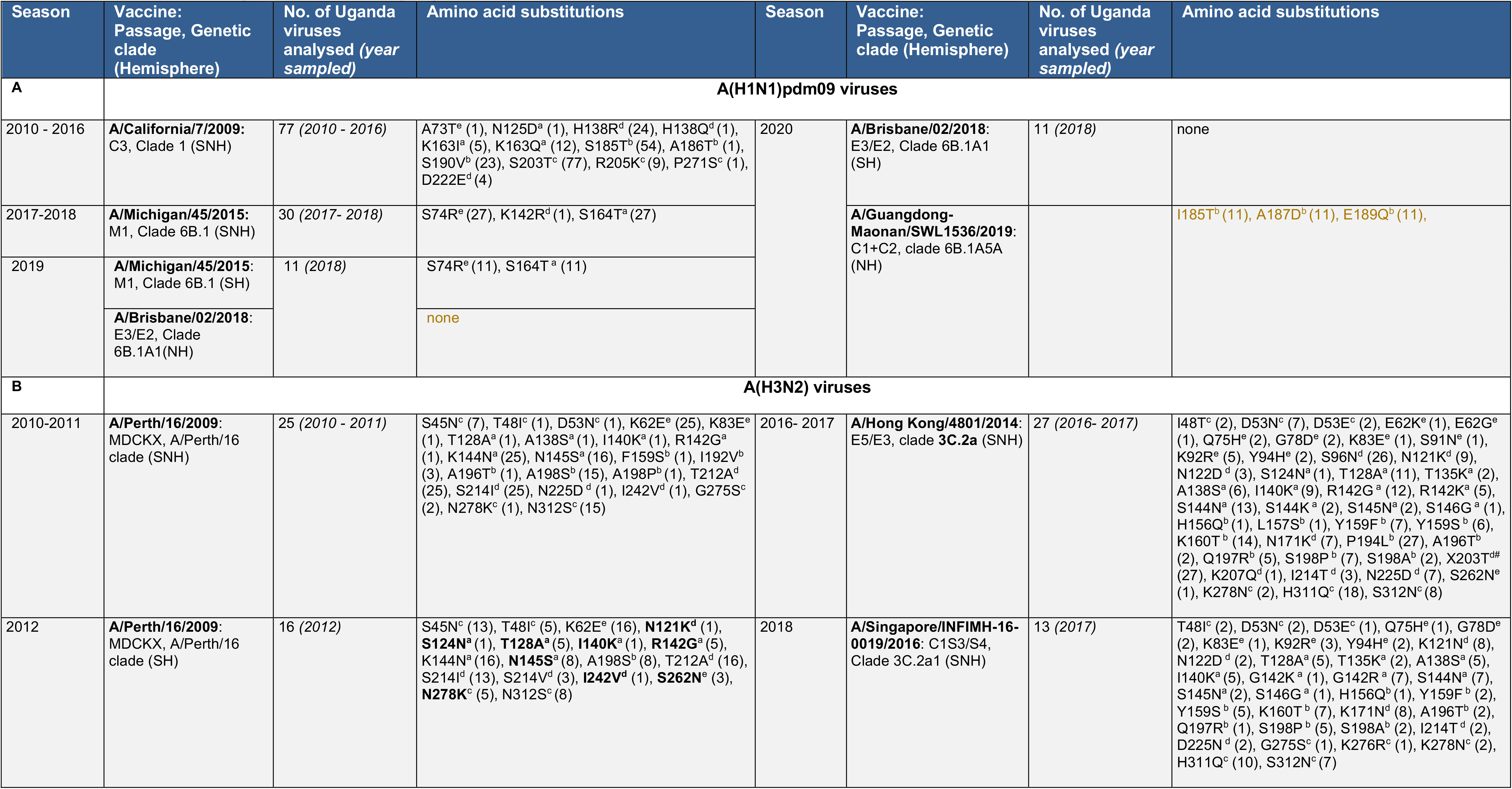
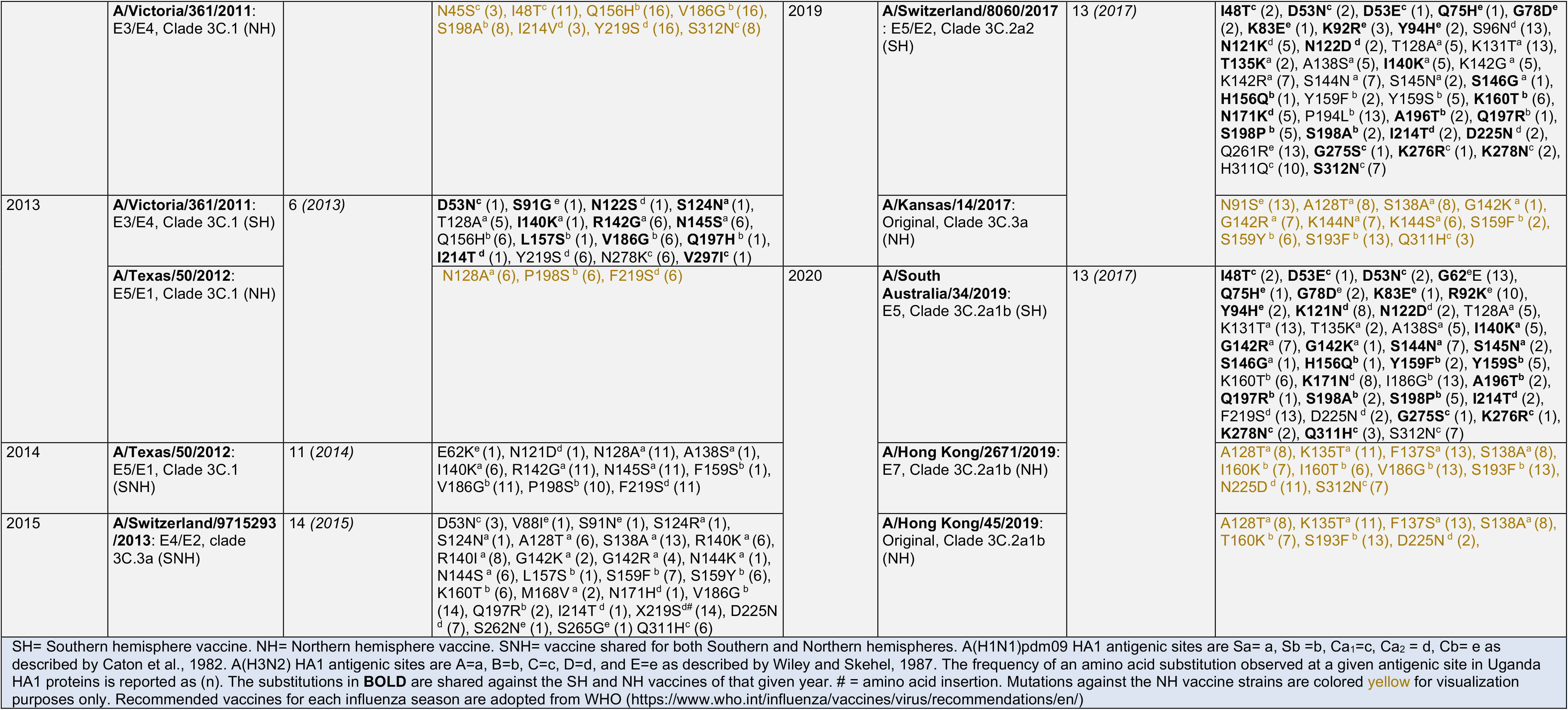
Antigenic drift among Uganda influenza A viruses. Mutations in the HA1 antigenic, receptor binding, and glycosylation sites of Uganda A(H1N1)pdm09 and A(H3N2) viruses compared to the Southern and Northern vaccine viruses.

| Season | Vaccine: Passage, Genetic clade (Hemisphere) | No. of Uganda viruses analysed (year sampled) | Amino acid substitutions | Season | Vaccine: Passage, Genetic clade (Hemisphere) | No. of Uganda viruses analysed (year sampled) | Amino acid substitutions |
| --- | --- | --- | --- | --- | --- | --- | --- |
| A | A(H1N1)pdm09 viruses |  |  |  |  |  |  |
| 2010 - 2016 | A/California/7/2009: C3, Clade 1 (SNH) | 77 (2010 - 2016) | A73T<sup>e</sup> (1), N125D<sup>a</sup> (1), H138R<sup>d</sup> (24), H138Q<sup>d</sup> (1), K163I<sup>a</sup> (5), K163Q<sup>a</sup> (12), S185T<sup>b</sup> (54), A186T<sup>b</sup> (1), S190V<sup>b</sup> (23), S203T<sup>c</sup> (77), R205K<sup>c</sup> (9), P271S<sup>c</sup> (1), D222E<sup>d</sup> (4) | 2020 | A/Brisbane/02/2018: E3/E2, Clade 6B.1A1 (SH) | 11 (2018) | none |
| 2017-2018 | A/Michigan/45/2015: M1, Clade 6B.1 (SNH) | 30 (2017- 2018) | S74R<sup>e</sup> (27), K142R<sup>d</sup> (1), S164T<sup>a</sup> (27) |  | A/Guangdong-Maonan/SWL1536/2019: C1+C2, clade 6B.1A5A (NH) |  | I185T<sup>b</sup> (11), A187D<sup>b</sup> (11), E189Q<sup>b</sup> (11), |
| 2019 | A/Michigan/45/2015: M1, Clade 6B.1 (SH) | 11 (2018) | S74R<sup>e</sup> (11), S164T<sup>a</sup> (11) |  |  |  |  |
|  | A/Brisbane/02/2018: E3/E2, Clade 6B.1A1(NH) |  | none |  |  |  |  |
| B | A(H3N2) viruses |  |  |  |  |  |  |
| 2010-2011 | A/Perth/16/2009: MDCKX, A/Perth/16 clade (SNH) | 25 (2010 - 2011) | S45N<sup>c</sup> (7), T48I<sup>c</sup> (1), D53N<sup>c</sup> (1), K62E<sup>e</sup> (25), K83E<sup>e</sup> (1), T128A<sup>a</sup> (1), A138S<sup>a</sup> (1), I140K<sup>a</sup> (1), R142G<sup>a</sup> (1), K144N<sup>a</sup> (25), N145S<sup>a</sup> (16), F159S<sup>b</sup> (1), I192V<sup>b</sup> (3), A196T<sup>b</sup> (1), A198S<sup>b</sup> (15), A198P<sup>b</sup> (1), T212A<sup>d</sup> (25), S214I<sup>d</sup> (25), N225D<sup>d</sup> (1), I242V<sup>d</sup> (1), G275S<sup>c</sup> (2), N278K<sup>c</sup> (1), N312S<sup>c</sup> (15) | 2016- 2017 | A/Hong Kong/4801/2014: E5/E3, clade 3C.2a (SNH) | 27 (2016- 2017) | I48T<sup>c</sup> (2), D53N<sup>c</sup> (7), D53E<sup>c</sup> (2), E62K<sup>e</sup> (1), E62G<sup>e</sup> (1), Q75H<sup>e</sup> (2), G78D<sup>e</sup> (2), K83E<sup>e</sup> (1), S91N<sup>e</sup> (1), K92R<sup>e</sup> (5), Y94H<sup>e</sup> (2), S96N<sup>d</sup> (26), N121K<sup>d</sup> (9), N122D<sup>d</sup> (3), S124N<sup>a</sup> (1), T128A<sup>a</sup> (11), T135K<sup>a</sup> (2), A138S<sup>a</sup> (6), I140K<sup>a</sup> (9), R142G<sup>a</sup> (12), R142K<sup>a</sup> (5), S144N<sup>a</sup> (13), S144K<sup>a</sup> (2), S145N<sup>a</sup> (2), S146G<sup>a</sup> (1), H156Q<sup>b</sup> (1), L157S<sup>b</sup> (1), Y159F<sup>b</sup> (7), Y159S<sup>b</sup> (6), K160T<sup>b</sup> (14), N171K<sup>d</sup> (7), P194L<sup>b</sup> (27), A196T<sup>b</sup> (2), Q197R<sup>b</sup> (5), S198P<sup>b</sup> (7), S198A<sup>b</sup> (2), X203T<sup>d#</sup> (27), K207Q<sup>d</sup> (1), I214T<sup>d</sup> (3), N225D<sup>d</sup> (7), S262N<sup>e</sup> (1), K278N<sup>c</sup> (2), H311Q<sup>c</sup> (18), S312N<sup>c</sup> (8) |
| 2012 | A/Perth/16/2009: MDCKX, A/Perth/16 clade (SH) | 16 (2012) | S45N<sup>c</sup> (13), T48I<sup>c</sup> (5), K62E<sup>e</sup> (16), N121K<sup>d</sup> (1), S124N<sup>a</sup> (1), T128A<sup>a</sup> (5), I140K<sup>a</sup> (1), R142G<sup>a</sup> (5), K144N<sup>a</sup> (16), N145S<sup>a</sup> (8), A198S<sup>b</sup> (8), T212A<sup>d</sup> (16), S214I<sup>d</sup> (13), S214V<sup>d</sup> (3), I242V<sup>d</sup> (1), S262N<sup>e</sup> (3), N278K<sup>c</sup> (5), N312S<sup>c</sup> (8) | 2018 | A/Singapore/INFIMH-16-0019/2016: C1S3/S4, Clade 3C.2a1 (SNH) | 13 (2017) | T48I<sup>c</sup> (2), D53N<sup>c</sup> (2), D53E<sup>c</sup> (1), Q75H<sup>e</sup> (1), G78D<sup>e</sup> (2), K83E<sup>e</sup> (1), K92R<sup>e</sup> (3), Y94H<sup>e</sup> (2), K121N<sup>d</sup> (8), N122D<sup>d</sup> (2), T128A<sup>a</sup> (5), T135K<sup>a</sup> (2), A138S<sup>a</sup> (5), I140K<sup>a</sup> (5), G142K<sup>a</sup> (1), G142R<sup>a</sup> (7), S144N<sup>a</sup> (7), S145N<sup>a</sup> (2), S146G<sup>a</sup> (1), H156Q<sup>b</sup> (1), Y159F<sup>b</sup> (2), Y159S<sup>b</sup> (5), K160T<sup>b</sup> (7), K171N<sup>d</sup> (8), A196T<sup>b</sup> (2), Q197R<sup>b</sup> (1), S198P<sup>b</sup> (5), S198A<sup>b</sup> (2), I214T<sup>d</sup> (2), D225N<sup>d</sup> (2), G275S<sup>c</sup> (1), K276R<sup>c</sup> (1), K278N<sup>c</sup> (2), H311Q<sup>c</sup> (10), S312N<sup>c</sup> (7) |
|  | **A/Victoria/361/2011:**<br>E3/E4, Clade 3C.1 (NH) |  | **N45S<sup>c</sup> (3), I48T<sup>c</sup> (11), Q156H<sup>b</sup> (16), V186G<sup>b</sup> (16), S198A<sup>b</sup> (8), I214V<sup>d</sup> (3), Y219S<sup>d</sup> (16), S312N<sup>c</sup> (8)** | 2019 | **A/Switzerland/8060/2017**<br>: E5/E2, Clade 3C.2a2 (SH) | 13 (2017) | **I48T<sup>c</sup> (2), D53N<sup>c</sup> (2), D53E<sup>c</sup> (1), Q75H<sup>e</sup> (1), G78D<sup>e</sup> (2), K83E<sup>e</sup> (1), K92R<sup>e</sup> (3), Y94H<sup>e</sup> (2), S96N<sup>d</sup> (13), N121K<sup>d</sup> (5), N122D<sup>d</sup> (2), T128A<sup>a</sup> (5), K131T<sup>a</sup> (13), T135K<sup>a</sup> (2), A138S<sup>a</sup> (5), I140K<sup>a</sup> (5), K142G<sup>a</sup> (5), K142R<sup>a</sup> (7), S144N<sup>a</sup> (7), S145N<sup>a</sup> (2), S146G<sup>a</sup> (1), H156Q<sup>b</sup> (1), Y159F<sup>b</sup> (2), Y159S<sup>b</sup> (5), K160T<sup>b</sup> (6), N171K<sup>d</sup> (5), P194L<sup>b</sup> (13), A196T<sup>b</sup> (2), Q197R<sup>b</sup> (1), S198P<sup>b</sup> (5), S198A<sup>b</sup> (2), I214T<sup>d</sup> (2), D225N<sup>d</sup> (2), Q261R<sup>e</sup> (13), G275S<sup>c</sup> (1), K276R<sup>c</sup> (1), K278N<sup>c</sup> (2), H311Q<sup>c</sup> (10), S312N<sup>c</sup> (7)** |
| 2013 | **A/Victoria/361/2011:**<br>E3/E4, Clade 3C.1 (SH) | 6 (2013) | **D53N<sup>c</sup> (1), S91G<sup>e</sup> (1), N122S<sup>d</sup> (1), S124N<sup>a</sup> (1), T128A<sup>a</sup> (5), I140K<sup>a</sup> (1), R142G<sup>a</sup> (6), N145S<sup>a</sup> (6), Q156H<sup>b</sup> (6), L157S<sup>b</sup> (1), V186G<sup>b</sup> (6), Q197H<sup>b</sup> (1), I214T<sup>d</sup> (1), Y219S<sup>d</sup> (6), N278K<sup>c</sup> (6), V297I<sup>c</sup> (1)** |  | **A/Kansas/14/2017:**<br>Original, Clade 3C.3a (NH) |  | **N91S<sup>e</sup> (13), A128T<sup>a</sup> (8), S138A<sup>a</sup> (8), G142K<sup>a</sup> (1), G142R<sup>a</sup> (7), K144N<sup>a</sup> (7), K144S<sup>a</sup> (6), S159F<sup>b</sup> (2), S159Y<sup>b</sup> (6), S193F<sup>b</sup> (13), Q311H<sup>c</sup> (3)** |
|  | **A/Texas/50/2012:**<br>E5/E1, Clade 3C.1 (NH) |  | **N128A<sup>a</sup> (6), P198S<sup>b</sup> (6), F219S<sup>d</sup> (6)** | 2020 | **A/South Australia/34/2019:**<br>E5, Clade 3C.2a1b (SH) | 13 (2017) | **I48T<sup>c</sup> (2), D53E<sup>c</sup> (1), D53N<sup>c</sup> (2), G62E<sup>e</sup> (13), Q75H<sup>e</sup> (1), G78D<sup>e</sup> (2), K83E<sup>e</sup> (1), R92K<sup>e</sup> (10), Y94H<sup>e</sup> (2), K121N<sup>d</sup> (8), N122D<sup>d</sup> (2), T128A<sup>a</sup> (5), K131T<sup>a</sup> (13), T135K<sup>a</sup> (2), A138S<sup>a</sup> (5), I140K<sup>a</sup> (5), G142R<sup>a</sup> (7), G142K<sup>a</sup> (1), S144N<sup>a</sup> (7), S145N<sup>a</sup> (2), S146G<sup>a</sup> (1), H156Q<sup>b</sup> (1), Y159F<sup>b</sup> (2), Y159S<sup>b</sup> (5), K160T<sup>b</sup> (6), K171N<sup>d</sup> (8), I186G<sup>b</sup> (13), A196T<sup>b</sup> (2), Q197R<sup>b</sup> (1), S198A<sup>b</sup> (2), S198P<sup>b</sup> (5), I214T<sup>d</sup> (2), F219S<sup>d</sup> (13), D225N<sup>d</sup> (2), G275S<sup>c</sup> (1), K276R<sup>c</sup> (1), K278N<sup>c</sup> (2), Q311H<sup>c</sup> (3), S312N<sup>c</sup> (7)** |
| 2014 | **A/Texas/50/2012:**<br>E5/E1, Clade 3C.1 (SNH) | 11 (2014) | **E62K<sup>e</sup> (1), N121D<sup>d</sup> (1), N128A<sup>a</sup> (11), A138S<sup>a</sup> (1), I140K<sup>a</sup> (6), R142G<sup>a</sup> (11), N145S<sup>a</sup> (11), F159S<sup>b</sup> (1), V186G<sup>b</sup> (11), P198S<sup>b</sup> (10), F219S<sup>d</sup> (11)** |  | **A/Hong Kong/2671/2019:**<br>E7, Clade 3C.2a1b (NH) |  | **A128T<sup>a</sup> (8), K135T<sup>a</sup> (11), F137S<sup>a</sup> (13), S138A<sup>a</sup> (8), I160K<sup>b</sup> (7), I160T<sup>b</sup> (6), V186G<sup>b</sup> (13), S193F<sup>b</sup> (13), N225D<sup>d</sup> (11), S312N<sup>c</sup> (7)** |
| 2015 | **A/Switzerland/9715293/2013:** E4/E2, clade 3C.3a (SNH) | 14 (2015) | **D53N<sup>c</sup> (3), V88I<sup>e</sup> (1), S91N<sup>e</sup> (1), S124R<sup>a</sup> (1), S124N<sup>a</sup> (1), A128T<sup>a</sup> (6), S138A<sup>a</sup> (13), R140K<sup>a</sup> (6), R140I<sup>a</sup> (8), G142K<sup>a</sup> (2), G142R<sup>a</sup> (4), N144K<sup>a</sup> (1), N144S<sup>a</sup> (6), L157S<sup>b</sup> (1), S159F<sup>b</sup> (7), S159Y<sup>b</sup> (6), K160T<sup>b</sup> (6), M168V<sup>a</sup> (2), N171H<sup>d</sup> (1), V186G<sup>b</sup> (14), Q197R<sup>b</sup> (2), I214T<sup>d</sup> (1), X219S<sup>d</sup> (14), D225N<sup>d</sup> (7), S262N<sup>e</sup> (1), S265G<sup>e</sup> (1), Q311H<sup>c</sup> (6)** |  | **A/Hong Kong/45/2019:**<br>Original, Clade 3C.2a1b (NH) |  | **A128T<sup>a</sup> (8), K135T<sup>a</sup> (11), F137S<sup>a</sup> (13), S138A<sup>a</sup> (8), T160K<sup>b</sup> (7), S193F<sup>b</sup> (13), D225N<sup>d</sup> (2),** |

SH= Southern hemisphere vaccine. NH= Northern hemisphere vaccine. SNH= vaccine shared for both Southern and Northern hemispheres. A(H1N1)pdm09 HA1 antigenic sites are Sa= a, Sb= b, Ca1= c, Ca2= d, Cb= e as described by Caton et al., 1982. A(H3N2) HA1 antigenic sites are A=a, B=b, C=c, D=d, and E=e as described by Wiley and Skehel, 1987. The frequency of an amino acid substitution observed at a given antigenic site in Uganda HA1 proteins is reported as (n). The substitutions in **BOLD** are shared against the SH and NH vaccines of that given year. # = amino acid insertion. Mutations against the NH vaccine strains are colored yellow for visualization purposes only. Recommended vaccines for each influenza season are adopted from WHO (https://www.who.int/influenza/vaccines/virus/recommendations/en/)

We observed 16 unique amino acid substitutions (UAAS) across the five antigenic sites amongst the 107 A(H1N1)pdm09 viruses (Table 1A). Ranking from the most variable site, Sa, Ca2, Sb, Ca1, and Cb had 4, 4, 3, 3, and 2 AAS, respectively. Substitution S164T, S185T, S203T, H138R, and S74R was most frequent at site Sa, Sb, Ca1, Ca2, and Cb, respectively. Seventy-percent (54/77) and 100% of 2010-2016 viruses carried S185T and S203T, respectively. Ninety-percent (27/30) of the 2017-2018 viruses carried S164T and S74R.

There were 93 UAAS across the five antigenic sites amongst the 99 A(H3N2) viruses (Table 1B). Ranked most variable was site A, D, B, C, and E with 24, 21, 20, 15, and 13 UAAS, respectively. Substitutions V186G and N145S were the most frequent in 47·47% (47/99) and 41·41% (41/99) of the all 2010-2017 A(H3N2) viruses. AAS at site A (K144N, N128A, R142G, and N145S), B (Q156H, V186G, P198S, and P194L), D (T212A, S214I, Y219S, and F219S), and E (K62E) were incorporated in all viruses analysed in a given season.

We observed mutated receptor binding sites (RBS) (A186T, S190V, and D222E) and glycosylation sites (GS) (A186T and N125D) of A(H1N1)pdm09 viruses previously described^18,22^. A(H3N2) viruses had more mutated RBS in the 130-loop (135, 138, and 140), 190-helix (186, 192, 194, and 196), 220-loop (219), and two GS (122 and 144) (Table 1B).

### Mutational landscape of complete HA, NA, and MP proteins

Uganda HA (H1), NA (N1), and MP proteins had 94, 81, and 21 UAAS, respectively, and were 97·85-99·09%, 98·04-98·97%, and 98·95-99·95% similar to A/California/7/2009, A/Michigan/45/2015, and A/Brisbane/02/2018 vaccines at amino acid level (sTable 4, Appendix pp 11-12).

N1 proteins lacked the neuraminidase inhibitors (NAIs) resistance substitution H275Y. However, 7·55% (8/106) N1 proteins had T362I (1), I117M (2), Y155H (2), and V234I (3) associated with reduced susceptibility to NAIs in vitro^22^.

Uganda HA (H3), NA (N2), and MP proteins had 161, 118, and 31 UAAS and amino acid similarity of 96·36-98·56%, 96·49-99·39%, and 97·48-99·86%, respectively, to A/Perth/16/2009, A/Victoria/361/2011, A/Texas/50/2012, A/Switzerland/9715293/2013, A/Hong Kong/4801/2014, A/Singapore/INFIMH-16-0019/2016, A/Switzerland/8060/2017, A/South Australia/34/2019, and A/Kansas/14/2017 vaccines (sTable 5, Appendix pp 13-16).

N2 proteins lacked the NAI-resistance H274Y (N2 numbering) but 17·8% (18/101) carried Y155F (2), E/D221D/K/E (15), and H374N (1) associated with reduced NAI susceptibility^22^.

All Uganda A(H1N1)pdm09 and A(H3N2) viruses incorporated adamantine-resistance marker (ARM) S31N. Seven-percent (8/104) of A(H3N2) M2 proteins carried a secondary ARM V27A relative to adamantine-susceptible A/New York/392/2004 virus (sFigure 5, Appendix pp 17).

### Temporal and spatial divergence of Uganda influenza A viruses

Uganda viruses phylogenetically clustered according to the year of collection with multiple lineages circulating annually (Figure 4). The A(H1N1)pdm09 viruses collected in 2010-2011 and 2012-2018 clustered distinctively. Two major lineages co-circulated in 2013 (Figure 4A). The H1 phylogeny showed lineage 1 (shaded area in blue) circulated in 2013-2016, while lineage 2 (shaded area in grey) persisted through 2018. In the N1 and MP phylogenies, lineage 1 and 2 emerged in 2013 and 2014, respectively, and co-circulated in 2014-2015. Lineage 2 dominated in 2016 and persisted through 2018. The 2017-2018 A(H1N1)pdm09 viruses monophyletically clustered at the tip in all phylogenies.

**Figure 4:**
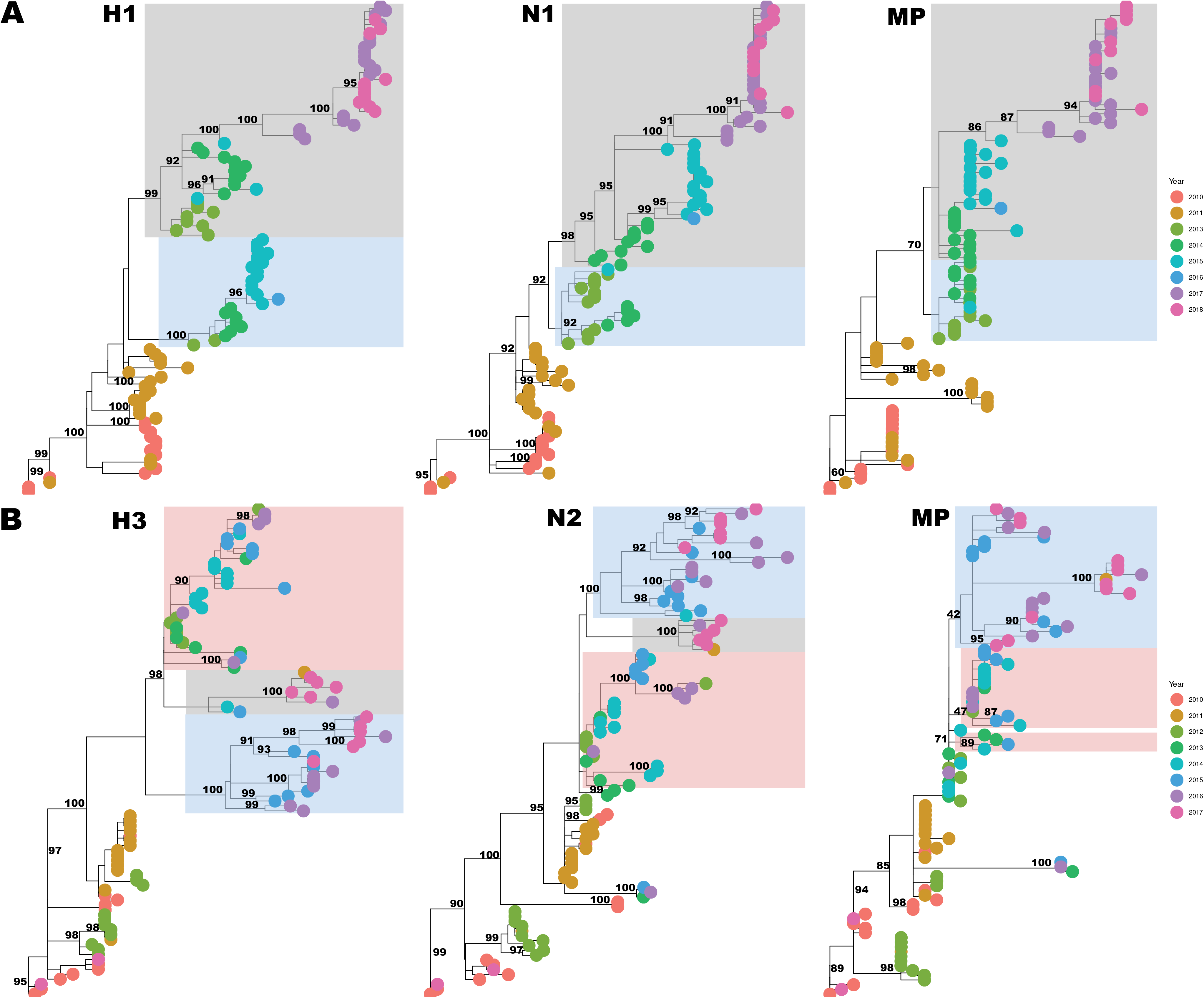
Phylogenies showing the temporal divergence of the HA, NA and MP genes of Uganda A(H1N1)pdm09 (A) and A(H3N2) (B) influenza viruses collected from 2010 to 2018. Trees were rooted using the oldest sequence in the Ugandan dataset. Shaded clusters are the two and three major co-circulating lineages observed since 2013 and 2012 for A(H1N1)pdm09 and A(H3N2) viruses, respectively. The third A(H3N2) lineage (with a 2011, 2016, 2017 viruses) disappeared in the MP gene tree.

One dominant A(H3N2) virus lineage (1) (blue) circulated in 2013-2016. Lineage 2 and 3 (pink and grey) emerged in 2015 and 2016, respectively, and co-circulated through 2017 (Figure 4B).

Viruses from different sites mixed in all phylogenies (sFigure 6, Appendix pp 18).

### Viral genetic clades circulating in Uganda

PhyCLIP grouped A(H1N1)pdm09 viruses in eight clusters, seven of which belonged to four ECDC global genetic clades (3, 5, 6, and 7) (Figure 5A). All four clades co-circulated in 2011. Clade 3 had 2010-2011 viruses carrying A134T and S183P substitutions. Clade 5 had 2011 viruses with D97N, R205K, I216V, and V249L. One 2011 virus from Kisenyi clustered with A/St.Petersburgh/100/2011 in clade 7 with A197T, S143G, and K163I. Clade 6 viruses with AAS D97N, S185T, and S203T dominated since 2012, diverging into 6A, 6B, and 6C. Clade 6B diverged into subclade 6B.1 with S162N, I216T, and S84N dominating in 2017-2018. A novel subclade 6B.1a with T120A emerged in November 2017 and circulated through 2018.

PhyCLIP grouped A(H3N2) viruses into 5 clusters belonging to global genetic clades 3 and 7 (Figure 5B). Clade 3 and 7 co-circulated in 2010-2012. Most viruses belonged to clade 3 carrying N145S and V223I. Clade 3 diverged into 3B, 3C, 3C.2a, and 3C.3. Subclade 3C.2a diverged into subclades 3C.2a1a with T135K (May-June 2017) and 3C.2a1b with K92R and H311Q (May-November 2017).

**Figure 5:**
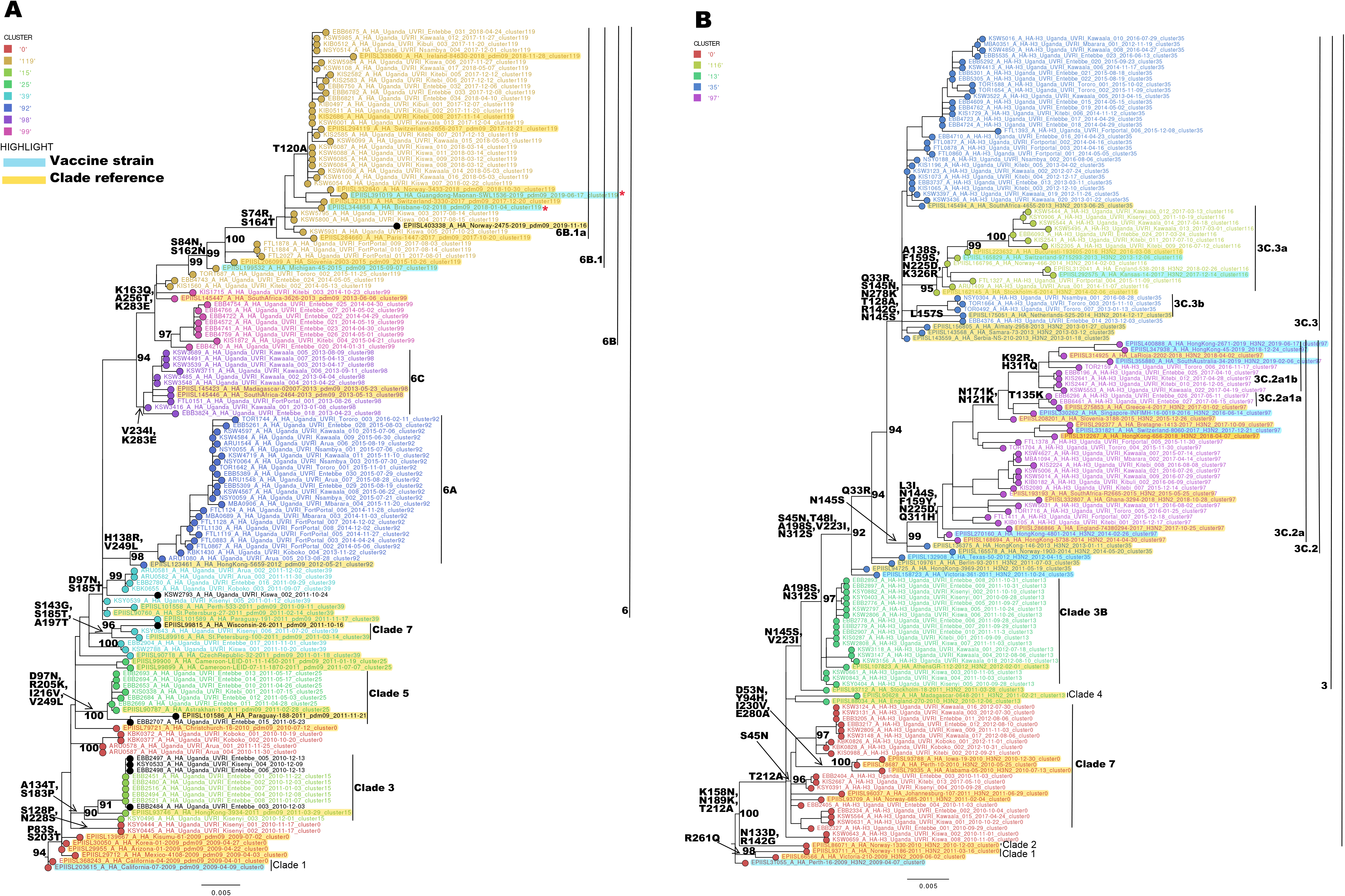
Genetic clades of influenza A viruses that previously circulated in 2010-2018 in Uganda. **Panel A** shows clusters and genetic clades for 2010-2018 A(H1N1)pdm09 viruses identified by statistical (PhyCLIP) and reference-based (ECDC) methods, respectively. Genetic clade 6 contains five PhyCLIP clusters (119, 92, 98, 99, and 39) with distinctive detection of 6A, 6B, and 6C. Clades 3, 5 and 7 are comparable to PhyCLIP clusters 15, 25, and 39, respectively. All clade 3, 5, and 7 viruses were collected from Entebbe and Kampala (Central Uganda) and circulated in 2010-2011. **Panel B** shows clusters and genetic clades for 2010-2017 A(H3N2) viruses identified by statistical (PhyCLIP) and reference-based (ECDC) methods, respectively. Genetic clade 3 contains four PhyCLIP clusters (13, 35, 97, and 116). Genetic clade 7 is comparable to PhyCLIP cluster 0. Clade 3 persisted in all the nine years while clade 7 re-emerged in 2017. The tip nodes and tip labels are coloured based on the cluster assigned by PhyCLIP, while labelled genetic clades (indicated by black bars) are those identified based on the amino acid mutations in the HA1 protein (ECDC method). Characteristic substitutions for each clade are indicated on the tree trunk in **bold**.

### Phylogenetic relatedness of African viruses

We define a group as a cluster with at least three Uganda sequences in the African gene tree.

Uganda A(H1N1)pdm09 viruses collected before 2016 clustered uniquely towards the root, while the 2017-2018 viruses mixed with others from Eastern, Central, Western, and Southern Africa (sFigure 7-9, Appendix pp 19-21). We observed 9 groups in each A(H1N1)pmd09 gene tree. Forty-four percent (4/9) of the H1 and N1 (2,4, 5, and 6), and MP (2, 4, 5, and 7) were unique to Uganda (sTable 6, Appendix pp 25).

Uganda A(H3N2) viruses collected in 2008-2016 and some before April 2017 clustered uniquely closest to the root (sFigure 10-12, Appendix pp 22-24). There were 10, 11, and 15 groups in the H3, N2, and MP gene trees, respectively. Two H3 (1 and 2), one N2 (6), and eight MP (1, 3, 5, 6, 8, 9, 12, and 13) groups had only Uganda viruses (sTable 6, Appendix pp 25). Notably, the Makerere Walter Reed project (MWRP)^8^ viruses collected in 2008-2009 clustered separately from our new viruses.

If not clustered alone, Uganda viruses interspersed with viruses from Eastern, Central, Western, and Southern Africa. A(H1N1)pdm09 viruses clustered uniquely with viruses from Seychelles, Burkina Faso, Sierra Leone, CoteD’iviore, Mozambique, and Egypt, while A(H3N2) viruses grouped uniquely with Reunion and Rwanda viruses. Interestingly, five unique H3 lineages co-circulated in 2019; one Kenyan and four in West Africa (not shown). Group details are described in sTable 6 (Appendix pp 25).

Accessions for sequences used in this study are provided in sTable 7 (Appendix pp 26-27).

## DISCUSSION

We have demonstrated the feasibility of NGS whole-genome sequencing of IAVs in low-income settings. Analysis of the generated sequences highlighted antigenic drift, multiple introductions, and local transmission of both A(H1N1)pdm09 and A(H3N2) influenza viruses in Uganda. All Uganda viruses carried the M2 adamantane-resistance marker S31N. Africa viruses were genetically similar but several unique viral lineages (with bootstrap >90) were observed in Uganda and other countries persisting for 1-3 years.

We successfully generated 96·58% (226/234) genomes and recovered 85·4% (193/226) WG directly from respiratory swabs. Our WG recovery rate is comparable to the 82-88% reported in developed countries^14,23^ and better than 47·3% and 66% reported in Scotland^24^ and Mexico^25^, respectively. 10-20% of frozen swabs fail NGS as the RNA quality gets compromised during collection, storage, and pre-sequencing analysis. The smallest genes, MP and NS, were sequenced at a greater depth and the least for the polymerase segment 2 (PB1) as previously reported in Mexico and France^23,25^.

Sequence analysis revealed several AAS in the HA, NA, and MP proteins of Uganda IAVs. Notably, antigenic sites Sa and Ca2 for A(H1N1)pdm09, and A, D and B for A(H3N2) viruses varied most. The A(H1N1)pdm09 virus substitutions S164T and S74R, also dominant the 2017-2018 Kenya viruses^26^. The frequent S203T at site Ca1 has been reported globally but its function is unknown^27^. We observed RBS D222E, speculated to increase infection severity, in 5·2% (4/77) of the 2010-2011 A(H1N1)pdm09 viruses^27^. H1 (P100S, S220T, I338V) and N1 (V106I, N248D) earlier detected in 2009-2011 in Uganda viruses against A/California/7/2009^9^ continued to circulate through 2017. All 2010-2013 Uganda A(H3N2) viruses had mutated antigenic sites also observed in Kenya^28^.

Uganda viruses lacked H274Y or H275Y confirming previous reports of 98% of global IAVs being sensitive to NAIs^6^. NAIs are not publicly available making their intake low in Uganda. However, some N1 proteins carried permissive AAS V234I, N369K, and V241I known to counteract the effect of H275Y. These AAS enhance the NA surface expression and enzymatic activity, hence increasing viral fitness. Uganda IAVs NA active and catalytic sites^29^ were highly conserved indicating lack of pharmacological selection pressure.

The A(H1N1)pdm09 viruses formed steeper ladder-like phylogenies than A(H3N2) viruses, indicating directional selection due to immune escape^30^. Further WG analysis to identify processes such as reassortment and selection pressures driving the observed genetic variations is ongoing.

Phylogenetic analysis also revealed the circulation of multiple viral lineages and global genetic clades in Uganda as observed worldwide^7,8,11^. New viral lineages and clades were observed in between seasons indicating multiple viral introductions into Uganda per year. Novel A(H3N2) virus subclades 3C.2a1a and 3C. 2a1b were introduced in 2017. Uganda viruses phylogenetically clustered with no geographical signal showing an extensive viral mixing and local transmissions.

Uganda A(H1N1)pdm09 and A(H3N2) viruses collected before 2015 and 2016, respectively, clustered distinctively from most Africa viruses. However, this could be due to insufficient representative viruses sequenced in earlier years across Africa. Viruses collected later interspersed with those from Eastern, Central, Southern, and Western Africa indicative of viral genetic similarity. Also, 2010-2018 Uganda viruses clustered with global viruses indicating that Uganda contributes to the African and global influenza ecology.

The coronavirus disease 2019 (COVID-19) pandemic revealed viral exchange among countries through borders, for example, Kenya and Uganda. Future studies should explore advanced phylodynamic and demographical models to determine key importers and migration rates of IAVs in Uganda.

This is the first and largest study to sequence Uganda A(H1N1)pdm09 and A(H3N2) virus WG locally, and internationally to describe a detailed temporal and spatial genetic diversity and evolution patterns of Africa IAVs. Our sequences add significantly to publicly available Africa data. However, we used pre-collected data with a biased geographical sampling. Therefore, we did not consider geographical location in the swab randomisation. We sequenced only 13 and 12 viruses from Eastern and Northwest, respectively. Due to financial constraints, we sampled sequenced only 24·1% (234/971) of available viruses, 8·12% (19/234) of which failed quality control and NGS. We did not include clinical data or do phenotypic analysis to assess the effect of the substitutions observed on viral virulence, pathogenicity, and transmissibility.

This study provides a platform for larger future studies and highlights the potential of genomics surveillance to improve viral detection and disease management. Existing African surveillance programmes should prioritize routine sequencing and genome analysis to monitor circulating IAVs. Our findings will inform Uganda’s public health, use of NAIs as a prophylactic treatment, and decision to formulate influenza vaccination programs, especially for high-risk groups like children, pregnant mothers, and the elderly.

## Authors’ contributions

GN proposed the idea and designed the study with supervision from SDWF, JMK, and DPK. NO, JTK and JJL manage the UVRI-NIC influenza surveillance and provided both the samples and patient epidemiological data. DCO, ZRD, CNA, and DJN hosted, provided protocol, and trained GN on laboratory and whole-genome sequencing procedures. GN analysed the data with guidance from SDWF, interpreted results, and wrote the manuscript. SDWF and DJ guided on result presentation. All authors had full access to all the data in the study and had final responsibility for the decision to submit for publication, and GN and SDWF verified the underlying data. All authors revised and edited the manuscript’s preceding drafts and approved the final manuscript.

## Data sharing

All sequences generated in this study with their respective metadata were submitted to GISAID EpiFlu (https://www.gisaid.org/) under accessions EPIISL498819 - EPIISL498931 [A(H1N1pdm09)], and EPIISL498934 - EPIISL499037 [A(H3N2)], and will be available publicly.

## Acknowledgments

We thank the participants for providing samples. We extend our gratitude to the effort and contribution of the surveillance clinical officers and nurses at the sentinel sites, and the laboratory team at the Department of Arbovirology Emerging & Re-Emerging Infectious Diseases (UVRI) in sample and data collection, processing, archiving, and permitting us to use them. The authors acknowledge the authors, originating and submitting laboratories of the GISAID influenza viral sequences analysed in this study. All data submitters can be contacted through the GISAID EpiFlu platform.

