## Supplementary Material for "Phylogenomic analysis of Uganda influenza type-A viruses to assess their relatedness to the vaccine strains and other Africa viruses: a molecular epidemiology study"

### **Supplementary materials and methods**

#### **Source of swabs and sampling**

Nasal and oropharyngeal swabs were collected from outpatients and inpatients with influenza-like illnesses (ILI) and severe acute respiratory illnesses (SARI) at the different sentinel sites, respectively, as described by Lutwama *et.al*^1^. The swabs were tested for influenza (A and B) and the IAV were further subtyped for seasonal [A(H1N1) and A(H3N2)] and pandemic A(H1N1)pdm09 influenza using the Centers for Disease Control’s (CDC) real-time reverse-transcription polymerase chain reaction (rRT-PCR) protocols and primers (Atlanta, Georgia)^2^ . All patient swabs were uniquely coded and frozen at -80^0^C, and their sociodemographic data recorded using EpiInfo (CDC, Atlanta)^1^.

#### **Viral RNA isolation and amplification**

Following isolation, the viral RNA was reverse transcribed into cDNA and the entire IAV genome amplified using the multi-segment real-time polymerase chain reaction (M-RTPCR)^3^ and universal IAV Uni/Inf primers in 25 μL reactions containing 8 μL nuclease-free water, 12.5 μL 2× RT-PCR buffer, 0.2 μL Uni12/Inf1 (10 μM), 0.3 μL Uni12/Inf3 (10 μM), 0.5 μL Uni13/Inf1 (10 μM), 0.5 μL SuperScript III One-Step RT-PCR with Platinum *Taq* High Fidelity (Invitrogen) and 3 μL extracted RNA. The M-RTPCR standardised thermocycling conditions were as follows: 42 °C for 50 minutes, 50 °C for 10 minutes, 94 °C for 2 minutes; 4 cycles (94 °C for 30 seconds, 43 °C for 30 seconds and 68 °C for 3 minutes and 50 seconds) followed by 30 cycles of 94 °C for 30 seconds, 57 °C for 30 seconds and 68 °C for 3 minutes and 30 seconds (with the 3 minutes and 30 seconds for the 68 °C extension step increased by 10 seconds per subsequent cycle after cycle 1); and a final extension step at 68 °C for 10 minutes.

#### **Next-generation sequencing**

Following PCR, the amplicons were purified using 1X AMPure XP beads (Beckman Coulter Inc., Brea, CA, USA), quantified with Quant-iT dsDNA High Sensitivity Assay (Invitrogen, Carlsbad, CA, USA), and normalized to 0.2 ng/μL. Indexed paired end libraries were then generated from 2.5 μL of 0.2 ng/μL amplicon pool using Nextera XT Sample Preparation Kit (Illumina, San Diego, CA, USA) following the manufacturer’s protocol. Amplified libraries were purified using 0.8X AMPure XP beads, quantitated with Quant-iT dsDNA High Sensitivity Assay (Invitrogen, Carlsbad, CA, USA), and evaluated for fragment size in the Agilent 2100 BioAnalyzer System using the Agilent High Sensitivity DNA Kit (Agilent Technologies, Santa Clara, CA, USA). Libraries were then diluted to 2nM in preparation for pooling and denaturation for running on the Illumina MiSeq (Illumina, San Diego, CA, USA). Pooled libraries were sodium hydroxide denatured, diluted to 12.5 pM and sequenced on the Illumina MiSeq using 2 x 250 bp paired end reads with the MiSeq v2 500 cycle kit (Illumina, San Diego, CA, USA). Five percent Phi-X (Illumina, San Diego, CA, USA) spike-in was added to the libraries to increase library diversity by creating a more diverse set of library clusters. The MiSeq generated paired reads as fastq.gz files for each sample.

#### **Sequence quality control**

Raw MiSeq reads were de-duplicated using FastUniq v1.1^4^ and Trimmomatic v0·39^5^ was used to trim off Nextera transposase, adaptors, and PCR primers from the unique reads, retaining only reads with ≥80 bps. Clean reads were used as input for the Iterative Refinement Meta-Assembler (IRMA) assembly.

#### **Genome assembly using Iterative Refinement Meta-Assembler (IRMA)**

IRMA default settings for IAV genome assembly were as follows: median read quality score (Q-score) filter of 30; minimum read length of 125; frequency threshold for insertion and deletion refinement of 0.25 and 0.6, respectively; mismatch penalty of 5; and gap opening penalty of 10^6^. The IRMA output included: consensus sequences for all the eight gene segments, paired read counts, coverage depth, allele frequencies, and statistically supported variants for each sample.

#### **Viral sequence clustering using the Phylogenetic Clustering by Linear Integer Programming (PhyCLIP)**

Maximum likelihood trees were rooted using the oldest sequence A/California/04/2009 for subtype A(H1N1)pdm09 and A/Perth/16/2009 for subtype A(H3N2). Rooted trees were used as input for PhyCLIP^7^. PhyCLIP clustering was optimised using a series of parameter sets for the minimum number of sequences [S, 3-10(1)], false discovery rate (*FDR*, 0.05-0.20(0.05)], multiple of deviations [*gamma*, 1-3(0.5)] and zero-branch length collapsed. The optimal clustered tree (with >99% of all sequences clustered) for subtype A(H1N1)pdm09 was obtained with S=7, FDR=0.2 and gamma = 3.0 , while for subtype A(H3N2) at S=10, FDR= 0.05 and gamma =3.0.

#### **sFigure 1**: **Geographical distribution of sampled sentinel sites in the general UVRI-NIC influenza surveillance programme**

**sFigure 1**: **Geographical distribution of sampled sentinel sites in the UVRI-NIC influenza surveillance programme**. District hospitals included Arua-ARU, Entebbe-EBB, Fort Portal-FTL, Koboko-KBK, Mbarara-MBA, Tororo-TOR, and Kampala. Kampala had three hospitals: International Hospital Kampala-IHK, Kibuli moslem hospital-KIB, Nsambya Hospital-NSY, and four clinics: Kawaala Health Center-KIS, Kiswa Health Center-KSW, Kisenyi Health Center-KSY, and Kitebi health centre-KIS samples. Swabs were assigned unique laboratory identification number using the site codes and a number. For example, EBB0001 for swab 0001 from Entebbe.

### **Supplementary results**

#### **sFigure 2: Swab sample selection for influenza whole-genome sequencing**

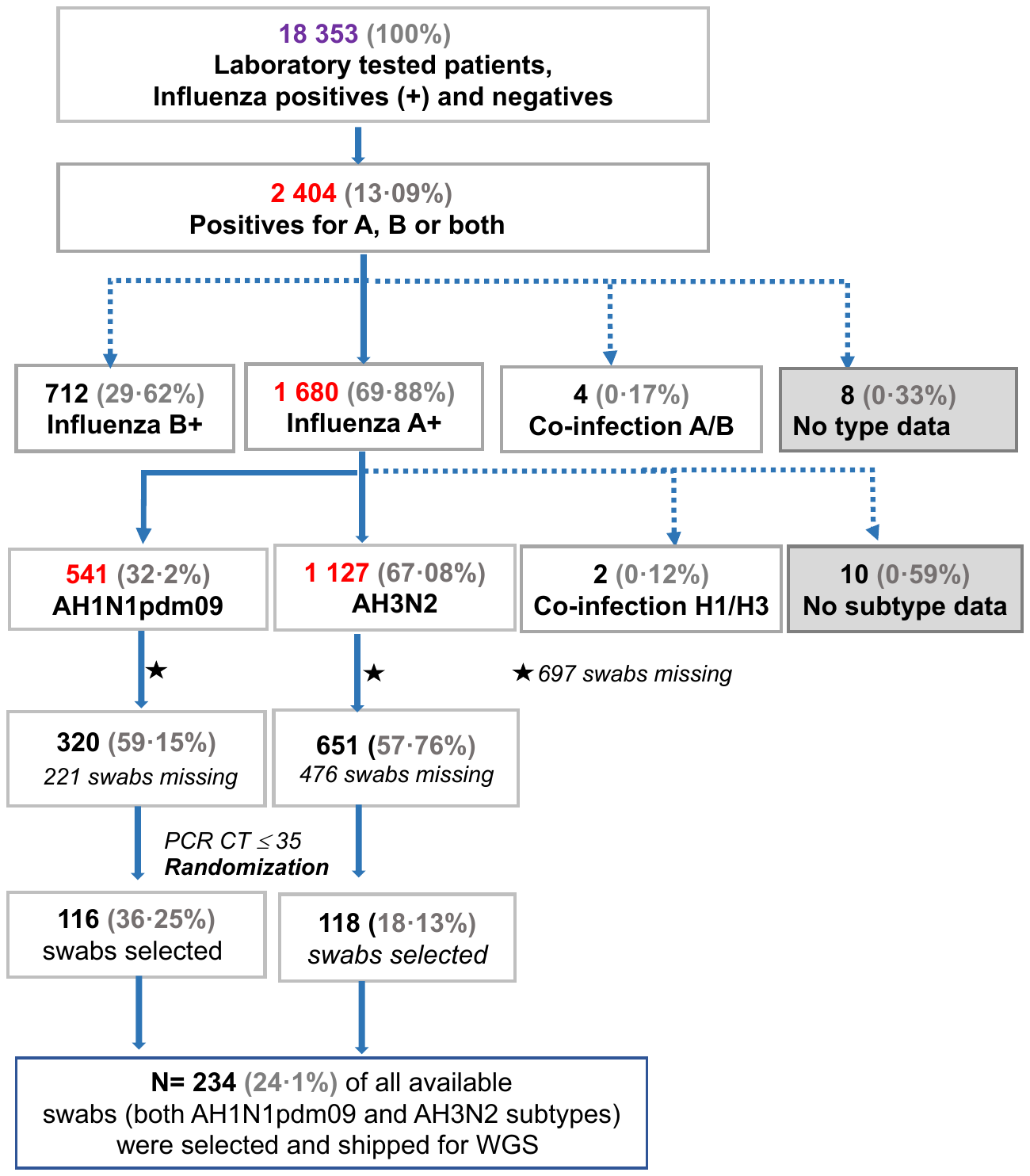

**SFigure 2: Flowchart showing how swabs were selected for influenza whole-genome sequencing (WGS).** Patients diagnosed with either influenza subtypes A(H1N1)pdm09 or A(H3N2) and whose swab had a PCR CT≤35 had their laboratory codes randomised based on the subtype and year of collection using the R software v3.6.3 (https://www.r-project.org). All available swabs were retrieved for years with less than fifteen swabs. In total, 24·1% (234/971) of the available swabs were selected for WGS. The 697 swabs missing include some shipped to the CDC for routine surveillance and some lost due to accidental failure of a freezer. The numbers are based on the UVRI-NIC laboratory dataset only, as of 9^th^ May 2018.

#### **sTable 1:** **Number of influenza A positive patients’ swabs that were available for sampling per subtype**

| **Year** | **Total number of swabs from A(H1N1)pdm09 positives** | **Number of swabs available for sampling (no PCR criteria)** | **Number of swabs available for sampling**  **PCR flu a subtype (CT≤35)** | **Number of swabs randomly sampled** | **Total number of swabs from A(H3N2) positives** | **Number of swabs available for sampling (no PCR criteria)** | **Number of swabs available for sampling**  **PCR flu a subtype (CT≤35)** | **Number of swabs randomly sampled** |
| --- | --- | --- | --- | --- | --- | --- | --- | --- |
|  | **A(H1N1)pdm09** | | | | **A(H3N2)** | | | |
| 2010 | 18 | 14 | 14 | 14 | 19 | 13 | 13 | 12 |
| 2011 | 93 | 71 | 67 | 19 | 51 | 45 | 44 | 16 |
| 2012 | 4 | 3 | 3 | 2 | 223 | 76 | 72 | 15 |
| 2013 | 71 | 23 | 22 | 15 | 30 | 19 | 19 | 15 |
| 2014 | 145 | 25 | 25 | 18 | 211 | 16 | 16 | 12 |
| 2015 | 144 | 130 | 130 | 17 | 118 | 100 | 100 | 14 |
| 2016 | 1 | 1 | 1 | 1 | 191 | 174 | 174 | 17 |
| 2017 | 51 | 42 | 42 | 19 | 284 | 208 | 206 | 17 |
| 2018 | 14 | 11 | 11 | 11 | 0 | 0 | 0 | 0 |
| **Total** | **541** | **320** | **315** | **116** | **1127** | **651** | **644** | **118** |

**sTable1:** **The number of IAV positive patient swabs that were available and randomly selected for WGS per subtype per year (2010-2018).** For years where the number of available swabs was less than 15, all available swabs were sampled. Following the first randomisation, 5 A(H1N1)pdm09 and 13 A(H3N2) swabs recorded as available could not be located in the storage freezers. A second randomisation (not based on year) was done excluding swabs from the first randomisation that had successfully been retrieved. The first 5 swabs for the A(H1N1)pmd09 and 13 swabs for the A(H3N2) were chosen bringing the total to 116 and 118, respectively. All numbers are based on the UVRI-NIC laboratory dataset sampled between 22^nd^ October 2010 and 9^th^ May 2018.

#### **sTable 2:** **Age distribution of influenza A positive patients sampled in the general UVRI-NIC surveillance programme and the study**

| **Age group** | **Num. of patients tested positive for influenza A** | **Num. of patients tested positive for H1N1** | **Num. of patients tested positive for A(H1N1)pdm09** | **Num. of patients tested positive for A(H3N2)** | **Num. of patients co-infected with**  **A(H3) and H1N1pdm09** | **Num. of IAV patients’ swabs sampled for WGS** |
| --- | --- | --- | --- | --- | --- | --- |
| 1 month-<2 years | 591 | 12 | 176 | 401 | 2 | 78 |
| 2-<5 years | 632 | 7 | 203 | 422 | 0 | 88 |
| 5-<15 years | 335 | 2 | 113 | 220 | 0 | 42 |
| 15-<50 years | 222 | 7 | 85 | 130 | 0 | 20 |
| 50-<65 years, | 15 | 1 | 6 | 8 | 0 | 2 |
| ≥ 65 years | 7 | 0 | 1 | 6 | 0 | 0 |
| **Total** | **1802** | **29** | **584** | **1187** | **2** | **230** |

**sTable 2:** **Number of patients that tested positive for influenza A [A(H1N1)pdm09 or A(H3N2)] per age group (years) recorded in the general UVRI-NIC surveillance programme dataset sampled from 24^th^ October 2007 to 17^th^ December 2018.** The extra two patients had a co-infection of AH1/AH3 and were excluded. The extra two patients had a co-infection of AH1/AH3 and were excluded bringing the total IAV positives to 1800. Four of the sampled swabs for sequencing lacked sociodemographic data.

#### **sFigure 3: Age comparison between the general UVRI-NIC surveillance programme, excluded, and sampled influenza A patients**

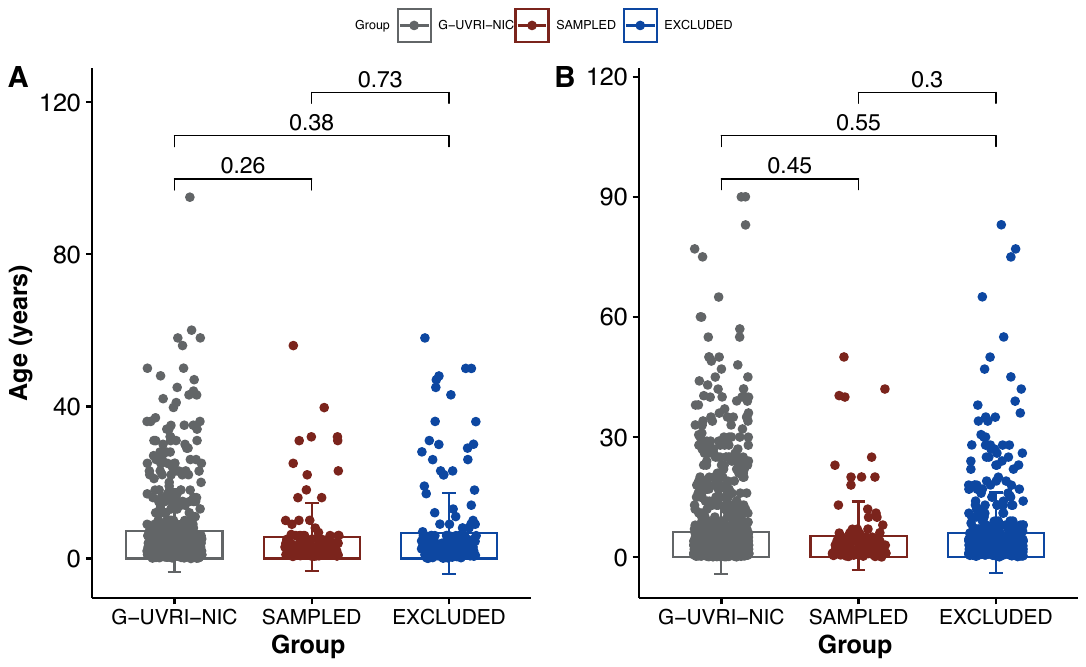
**sFigure 3: Age comparison between the general UVRI-NIC surveillance programme, excluded, and sampled influenza A patients.** Group definitions: G-UVRI-NIC are all IAV patients recorded in general UVRI-NIC surveillance database (N=1800), SAMPLED are IAV patients whose swabs were sampled in this study, and EXCLUDED are IAV patients whose swabs were not sequenced due to financial constraints. Group comparisons were done using the Wilcoxon test at p-value threshold 0·05. **Panel A** shows ages for A(H1N1)pdm09 patients and **Panel B** shows ages for A(H3N2) patients. 20 excluded patients’ swabs (9 H1N1pdm09 and 11 H3N2) had no sociodemographic data.

#### **sFigure 4: Comparison of the number of ILI and SARI cases and males and females between the general UVRI-NIC surveillance programme, excluded, and sampled influenza A patients**

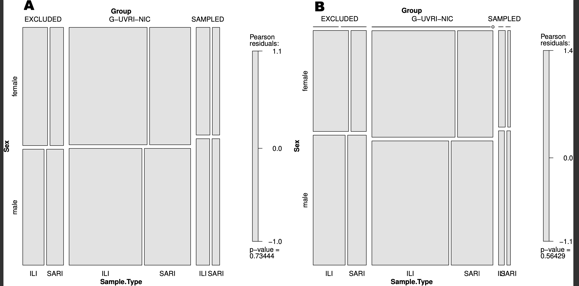

**sFigure 4: Mosaic graph showing the number of ILI and SARI cases and males and females between the general UVRI-NIC surveillance programme, excluded, and sampled influenza A patients.** **Sample.Type** correspond to the cases [influenza like illness (ILI) and severe acute respiratory illnesses (SARI). Group definitions as above in sFigure 3. Group comparisons were done using the chi-square test at p-value threshold 0·05. **Panel A** shows differences in group cases and gender for A(H1N1)pdm09 patients with p-value = 0·73 and **Panel B** shows differences in group cases and gender for A(H3N2) patients with p-value 0.56. 20 excluded patients’ swabs (9 H1N1pdm09 and 11 H3N2) had no sociodemographic data.

#### **sTable 3:** **Number of swabs from influenza A positive patients tested per site per year**

| **Region** | **Site** | **Number of swabs from A(H1N1)pdm09 positives** | **Year (number of swabs sampled that year)** | **Number of swabs from A(H3N2) positives** | **Year (number of swabs sampled that year)** |
| --- | --- | --- | --- | --- | --- |
|  |  | **A(H1N1)pdm09** | | **A(H3N2)** | |
| Northwest | Arua | 24 | **2010** (4); **2011** (3); **2012** (2); **2013** (3); **2014** (2); **2015** (10); **2016** (0); **2017** (0); **2018** (0) | 34 | **2010** (1); **2011** (0); **2012** (10); **2013** (0); **2014** (12); **2015** (2); **2016** (8); **2017** (1); **2018** (0) |
| Central | Entebbe | 130 | 2010 (7); 2011 (33); 2012 (0); 2013 (20); 2014 (29); 2015 (31); 2016 (0); 2017 (6); 2018 (4) | 244 | **2010** (10); **2011** (16); **2012** (58); **2013** (8); **2014** (52); **2015** (13); **2016** (28); **2017** (59); **2018** (0) |
| Western | Fort Portal | 47 | **2010** (0); **2011** (0); **2012** (0); **2013** (2); **2014** (26); **2015** (8); **2016** (0); **2017** (11); **2018** (0) | 69 | **2010** (0); **2011** (0); **2012** (0); **2013** (1); **2014** (17); **2015** (17); **2016** (26); **2017** (8); **2018** (0) |
| Northwest | Koboko | 21 | 2010 (2); 2011 (4); 2012 (0); 2013 (7); 2014 (1); 2015 (7); 2016 (0); 2017 (0); 2018 (0) | 51 | **2010** (0); **2011** (1); **2012** (5); **2013** (6); **2014** (22); **2015** (0); **2016** (7); **2017** (10); **2018** (0) |
| Western | Mbarara | 26 | **2010** (0); **2011** (8); **2012** (0); **2013** (3); **2014** (7); **2015** (8); **2016** (0); **2017** (0); **2018** (0) | 37 | **2010** (0); **2011** (0); **2012** (7); **2013** (2); **2014** (7); **2015** (2); **2016** (3); **2017** (16); **2018** (0) |
| Eastern | Tororo | 42 | **2010** (0); **2011** (1); **2012** (0); **2013** (1); **2014** (35); **2015** (4); **2016** (1); **2017** (0); **2018** (0) | 96 | **2010** (0); **2011** (0); **2012** (18); **2013** (2); **2014** (11); **2015** (36); **2016** (7); **2017** (22); **2018** (0) |
| Central | International Hospital Kampala-IHK | 1 | **2010** (0); **2011** (0); **2012** (0); **2013** (0); **2014** (0); **2015** (1); **2016** (0); **2017** (0); **2018** (0) | 10 | **2010** (0); **2011** (0); **2012** (0); **2013** (0); **2014** (0); **2015** (0); **2016** (10); **2017** (0); **2018** (0) |
| Central | Kibuli (KIB) | 9 | **2010** (0); **2011** (0); **2012** (0); **2013** (0); **2014** (0); **2015** (1); **2016** (0); **2017** (8); **2018** (0) | 27 | **2010** (0); **2011** (0); **2012** (0); **2013** (0); **2014** (0); **2015** (4); **2016** (6); **2017** (17); **2018** (0) |
| Central | Nsambya | 19 | **2010** (0); **2011** (0); **2012** (0); **2013** (0); **2014** (0); **2015** (13); **2016** ()0; **2017** (6); **2018** (0) | 24 | **2010** (0); **2011** (0); **2012** (0); **2013** (0); **2014** (0); **2015** (6); **2016** (5); **2017** (13); **2018** (0) |
| Central | Kawaala & Kitebi (KIS) | 73 | **2010** (0); **2011** (12); **2012** (0); **2013** (15); **2014** (13); **2015** (23); **2016** (0); **2017** (10); **2018** (0) | 197 | **2010** (0); **2011** (5); **2012** (60); **2013** (7); **2014** (34); **2015** (20); **2016** (28); **2017** (43); **2018** (0) |
| Central | Kiswa (KSW) | 129 | **2010** (0); **2011** (17); **2012** (2); **2013** (20); **2014** (32); **2015** (38); **2016** (0); **2017** (10); **2018** (10) | 324 | **2010** (5); **2011** (20); **2012** (65); **2013** (3); **2014** (56); **2015** (18); **2016** (62); **2017** (95); **2018** (0) |
| Central | Kisenyi (KSY) | 20 | **2010** (5); **2011** (15); **2012** (0); **2013** (0); **2014** (0); **2015** (0); **2016** (0); **2017** (0); **2018** (0) | 13 | **2010** (3); **2011** (9); **2012** (0); **2013** (1); **2014** (0); **2015** (0); **2016** (0); **2017** (0); **2018** (0) |
|  | No site data |  |  | 1 |  |
|  | Total | **541** |  | **1127** |  |

**sTable 3:** **Number of swabs from patients that tested positive for influenza A [A(H1N1)pdm09 or A(H3N2)] per site per year.** All numbers are based on the UVRI-NIC laboratory dataset sampled between 22^nd^ October 2010 and 9^th^ May 2018.

#### **sTable 4: Amino acid substitutions (AASs) in Uganda A(H1N1)pdm09 viruses compared to Southern and Northern Hemisphere vaccines viruses**

| Season | Vaccine:  Passage, Genetic clade (Hemisphere) | No. of Uganda viruses analysed *(year sampled)* | Total unique AAS; Average AAS per gene | Average AA similarity (%) | Virulence | Strong/Mild drug resistance | Host specificity shift | Antigenic drift/ Escape mutant | Creates/Removes potential  glycosylation site | Structural interactions | Other |
| --- | --- | --- | --- | --- | --- | --- | --- | --- | --- | --- | --- |
| Hemagglutinin (H1) | | | | | | | | | | | |
| 2010 - 2016 | **A/California/7/2009:**  C3, Clade 1 (SNH) | 77 *(2010 - 2016)* | 78; 11 | 97.85 | **E391K:61**  (14,886) | **P100S^+^:76**  (24,202), | **R269K:1**(7), **A151T:10**(259)  **D239E:4**(669), | **S200P:10**(4,314)**, N142D:1**(382), **S145P:3**(376), **S207V:23**(1),  **K180I:5(122),**  **K180Q:12**(1,022) |  | **N458K:1, A13T:1, A203T:1,A214T:1, A273T:12, A90T:1, D103E:1 D103N:1, D114N:58, D103N:1, D114N:58, D518E:1, D52N:1, E373A:1, E516K:58, E516Q:1, F12L:1, G56R:1, H143Y:2, H155Q:1, H155R:24 H290Y:1, I166M:1, I233V:7, I284T:4 I303L:1, I303V:7 I303X:1, I341V:2, I392V:1, I435V:2 I517V:1, I74T:1 K180I:17, K180Q:12 K300E:44,K419R:2, K475E:1, K475R:1 M274V:1, N146D:1, N245S:2, N458K:1, P135X:1, P199Q:1 P288S:1, R222K:9, R62K:1, S101I:1 S101N:1, S160G:2 S202T:54, S468N:54 S86P:2, T137A:1 T258K:3, V251I:10, V289A:2, V289I:15, V36I:3, V444I:2, V47X:1, V64I:** | **I338V^@^:75**(22,922**), P100S^@^:76**(24,062)  **S220T^@^:77**(22,514)**, I533V:1, L58I:1, L8M:1, R526K:23, V266L:33, V537A:24, V6A:1** |
| 2017-2018 | **A/Michigan/45/2015:**  E3, Clade 6B.1 (SNH) | 30 *(2017- 2018)* | 15; 5 | 99·09 |  |  | **R240Q^+^:30**  (8,199) | **K53N^+^:1**(10), **K159R:1**(40),  **R240Q:30**  (8,199), | **T295X^R^:1*** | **A278D:1, I113V:3, I303V:1, I312V:27, K516E:1, N472S:1, S181T:11, S91R:27, T137A:23, V36I:1,** | **R526G:1** |
| 2019 | **A/Michigan/45/2015**:  E3, Clade 6B.1 (SH) | 11 *(2018)* | 8; 5 | 99.07 |  |  | **R240Q^+^:11**  (8,199) | **R240Q:11**  (8,199) | **T295X^R^:1*** | **I303V:1, I312V:11, N472S:1, S181T:11, S91R:11, T137A:10** |  |
|  | **A/Brisbane/02/2018**:  E3, Clade 6B.1A1 (NH) |  | 9;6 | 98·9 |  |  | **R240Q^+^:11**  (8,199) | **P200S:11**(501), **R240Q:11**  (8,199 | **T295X^R^:1*** | **A299P:11, G62R:11, I303V:1, N472S:1, T137A:10, V315I:11** |  |
| Neuraminidase (N1) | | | | | | | | | | | |
| 2010-2016 | A/California/7/2009;  Clade1 (NH & SH) EPIISL203615 | 2010 - 2016 (n=76) | 62; 7 | 98.04 | **I365T:1**(335) | **Y155H^S^:2**(63),  **V267A^S+^:1(**13),  **I117M^M^:2**(325), **N248D^+^:75**(17,494), **N248X:1***, **V106I^+^:32**(9,341) |  | **I365T:1**(335),  **A343T:1**(4), **N200S:44**(8,216), **K432E:27 (**7,632**), N369K:59**(10,324), | **A343T:1**(4), **N44S^C^:45**  (8,932), **P272T^C^:2**(2), **N386K^R^:23**(7,307) | **A76T:1, A86V:1, C49Y:1, F115L:1 F74S:1, G382R:2, G454S:1, I321V:27, I389K, I389T:1, K262R:2, K84E:1, K84R:5, L85I:1, N141S:1, N270K:1, N397K:3, P93H:1, Q313R:9, R220K:9, S299A:1, S339L:2 S339P:1, T452I:5, V166I:14, V264I:1, V394I:27, V448L:1, V453X:1, Y353X:1** | **N248D^@^:75**(17,352), **V106I^@^:32**(9,340), **A20T:1, A20V:5, E47G:1, I314M:5, I34V:27, I359M:1, I46T:1, I46V:1, L40I:24, N42D:1, N71X:1, Q43K:2, Q45H:2, S442G:1, S82P:3, T9I:2, V13I:1, V241I:59** |
| 2017-2018 | **A/Michigan/45/2015:**  E3, Clade 6B.1 (SNH) | 2017-2018 (n=30) | 20; 6 | 98·47 | **I365T:23**(186) |  |  | **I365T:23**(186), **S366I:1**(4)**, I396V:1**(26)**,** |  | **A75V:1, D416N:24, G454D:1, G77R:27, I108M:1, I188T:27, K260E:1, N449D:27, P93H:1, L412S:1, S366I:1, T72I:24, V234I:3** | **Q45P:1, T16A:1, T362I:1, V80M:1, V81A:27** |
| 2019 | **A/Michigan/45/2015**:  E3, Clade 6B.1 (SH) | 2018 (n=11) | 16; 7 | 98·34 |  |  |  | **I365T:10**(186), **S366I:1**(4)**,** |  | **A75V:1, D416N:11, G454D:1, G77R:11, I108M:1, I188T:11, K260E:1, N449D:11, S366I:1, T72I:11** | **Q45P:1, T16A:1, T362I:1, V80M:1, V81A:11** |
|  | **A/Brisbane/02/2018**:  E3, Clade 6B.1A1 (NH) |  | 13; 4 | 98·97 |  |  |  | **I365T:10**(186), **S366I:1**(4)**,** |  | **A75V:1, D416N:11, G454D:1, I108M:1 K260E:1, S366I:1, T72I:11** | **Q45P:1, T13I:11, T16A:1, T362I:1, V80M:1** |
| Matrix Protein (M1 and M2) | | | | | | | | | | | |
| 2010-2016 | A/California/7/2009;  Clade1 (NH & SH) EPIISL203615 | 2010 - 2016 (n=79) | 16; 1·6 | 98·95 |  |  | **N133S:1**(43), **E16G:1**(52),  **G89D:1**(14) |  |  | **A33T:1, D21G:50, D21V:2** | **A227T:1, K230R:45, M192V:45, Q208K:1, T167A:1, V80I:54, E14G:4, R18K:6, T11I:1, Y76F:1** |
| 2017-2018 | **A/Michigan/45/2015:**  E3, Clade 6B.1 (SNH) | 2017-2018 (n=30) | 4; 1 | 99·86 |  |  |  |  |  | **F55L:1, I28F:3** | **E204X:1, M203I:1** |
| 2019 | **A/Michigan/45/2015**:  E3, Clade 6B.1 (SH) | 2018 (n=11) | 1; 1 | 99·95 |  |  |  |  |  | **F55L:1** |  |
|  | **A/Brisbane/02/2018**:  E3, Clade 6B.1A1 (NH) | 2018 (n=11) | 2; 1 | 99·75 |  |  |  |  |  | **F55L:1** | **X167T:11** |
| Data are in n or n (%), unless otherwise indicated. AA= Amino acid. AAS= Amino acid substitutions. SH= Southern hemisphere vaccine. HN= Northern hemisphere vaccine. SNH= vaccine strain shared by both the Southern and Northern hemispheres for a given influenza season. The number of unique amino acid substitutions observed in all proteins sequences and per protein sequence are represented as N; n. The number of times a substitution is observed in the Uganda virus proteins and globally is reported in BOLD and (bracket), respectively. Substitutions are colour-coded based on their known or predicted biological function and level of significance. Red mutations are the most significant (interestlevel =3) as they alter virulence, cause strong drug resistance and reverse premature stop codon in PB1-F2. The Orange (significant, interestlevel=2) occur at drug binding sites, affect host specificity and cause antigenic shift and mild drug resistance. Magenta (significant, interestlevel=2) adds or removes glycosylation sites. Blue mutations (moderately significant, interestlevel=1) have structural functions at interaction sites. Structural functions include host cell receptor binding, binding small ligand(s), viral oligomerization interfaces, and antibody recognition sites. The Green mutations (least significant, interestlevel=0) are common to subtypes while Black (least significant with warnlevel=0) have no known effects. Superscript symbol definitions: “*” = AAS reported for the first time (current global frequency = 0). “^+^” = AAS have their function reported in combination with others. “^R^” = AAS removes a potential glycosylation site. “^C^” = AAS creates a potential glycosylation site. “^S^” = AAS causes strong drug resistance. “^M^” = AAS causes mild drug resistance. “^@^” = AAS is a common subtype marker. | | | | | | | | | | | |

**STable 4: Amino acid substitutions in the complete coding sequences of HA (H1), NA (N1) and MP (M1 and M2) of Uganda A(H1N1)pdm09 viruses compared to Southern and Northern Hemisphere vaccines**. Amino acid similarity between protein sequences, amino acid substitutions and their corresponding global frequencies and functions were obtained from Flusurver (<http://flusurver.bii.a-star.edu.sg>; accessed on 30^th^ March 2021). The highest number of unique mutations in the H1 was 78, 62 in N1, and 16 in MP observed in 2010-2016 viruses against the A/California/7/2009 vaccine strain.

#### **sTable 5: Amino acid substitutions (AASs) in Uganda A(H3N2) viruses compared to Southern and Northern Hemisphere vaccines viruses**

| Season | Vaccine:  Passage, Genetic clade (Hemisphere) | No. of Uganda viruses analysed *(year sampled)* | Total unique AAS; Average AAS per gene | Average AA similarity (%) | Virulence &  Host specificity shift | Virulence & Antigenic drift / Escape mutant | Antigenic drift / Escape mutant | Strong/Mild drug resistance | Host specificity shift | Creates/Removes potential glycosylation site | Structural interactions | Other |
| --- | --- | --- | --- | --- | --- | --- | --- | --- | --- | --- | --- | --- |
| Hemagglutinin (H3) | | | | | | | | | | |  |  |
| 2010-2011 | **A/Perth/16/2009**: MDCKX, A/Perth/16 clade (SNH) | 25 *(2010 – 2011)* | 37; 9 | 98·13 | **A154S:1**  (3,152) | **A214S:15**(20,232), **A214P:1**(769), **N161S:24** (205,17) | **I156K^+^:1**(97), **K160N:25**(8,203), **R158G:1**(7,817), **K78E:25** (21,726), |  | **S230I:25**(24,366), **K99E^+^:1** (168), **A212T:1**(6), **I208V:3**(53),  **T144A^+^:1** (5,294), **F175S:1**(2,979),  **N241D:1** (16,616), | **K160N^C^:25**(8,203), **T144A^R^:1**(5,294), **S61N^C^:7**(20,569), | **A492T:3, D503N:15, Y9C:1, D505N:2, D69N:1, G291S:2, I258V:1, I422V:2 I434V:1, K280R:1, K2M:1, L199H:25 N294K:1, N328S:15, T64I:1, P305S:2, Q49R:1, R285K:2, T228A:25, V239I:16** | **I538M:1** |
| 2012 | **A/Perth/16/2009**: MDCKX, A/Perth/16 clade (SH) | 16 *(2012)* | 28; 11 | 97·83 |  | **A214S:8**(20,232), **N161S:8** (205,17) | **I156K^+^:1**(97),  **K160N:16**(8,203), **R158G:5**(7,817), **K78E:16** (21,726), |  | **S230I:13**(24,366), **T144A^+^:5** (5,294), **S230V:3** (30), | **K160N^C^:16**(8,203), **S140N^R^:1**(145), **S61N^C^:13**(20,569), **T144A^R^:5** (5,294) | **D503N:3, D505N:2, Y9C:5**  **I258V:1, K280R:8, T64I:5, L199H:16, L19V:1, N137K:1, N294K:5, N328S:8, Q49R:5, S25R:1, S278N:3, T228A:16, V14G:1, V239I:8,** |  |
|  | **A/Victoria/361/2011**: Clade 3C.1 (NH) |  | 27; 9 | 98·26 |  | **S214A:8**(778), **N161S:8** (20,312) | **I156K^+^:1**(90),  **Q172H:16**(21,478), **V202G:16**(21,412), **R158G:5**(7,817), |  | **I230V:3** (26), | **N61S^R^:3**(1230), **S140N^R^:1**(126), **T144A^R^:5** (5,288) | **D503N:3, D505N:2, H9C:5, H9Y:11, I239V:9, I64T:11, K280R:8, L19V:1, N137K:1, N294K:5, Q49R:5, S25R:1, S278N:3, S328N:11, V14G:1, Y235S:16** |  |
| 2013 | **A/Victoria/361/2011**: Clade 3C.1 (SH) | 6 *(2013)* | 21; 11 | 97·99 |  | **L173S:1**(710), **N161S:8** (20,312) | **I156K^+^:1**(90),  **Q172H:6**(21,478), **V202G:6**(21,412),  **N138S:1**(31),  **R158G:6**(7,721) | **S107G^M^:1**(11),  **N138S^M^:1**(31) | **I230T:1**(200),  **Q213H:1**(240), | **S140N^R^:1**(126), **T144A^R^:6** (5,288) | **D69N:1, H9Y:6, I422V:1, L19V:1, N294K:6, Q49R:6, V313I:1, V363M:1, Y235S:6** |  |
|  | **A/Texas/50/2012**: E5, Clade 3C.1 (NH) |  | 18; 8 | 98·49 |  | **P214S:6**(19,232), **L173S:1**(709),  **N161S:6**(19,737) | **V202G:6** (20,336), **I156K^+^:1**(90),  **N138S:1**(30), **R158G:6**(7,663) | **S107G^M^:1**(10), **N138S^M^:1**(30) | **I230T:1**(197),  **Q213H:1**(240),  **T144A^+^:6**(5,281), | **N138S^R^:1**(30), **S140N^R^:1**(122), | **D69N:1, F235S:6, I422V:1, L19V:1, V313I:1 V363M:1** |  |
| 2014 | **A/Texas/50/2012**: E5, Clade 3C.1 (SNH) | 11 (*2014)* | 17; 8 | 98·56 | **A154S:1**  (3,132), | **P214S:10**(19,232)**, N161S:11**(19,737), | **I156K^+^:1**(90), **V202G:11**(20,336),  **E78K:1**(562), **R158G:11**(7,663) |  | **F175S:1(**2,973),  **N241D:1** (16,534), **N144A^+^:11**(5,281) |  | **A16T:3, F235S:11 I10V:1 K342R:1, L19I:1, L19V:6, N137D:1** |  |
| 2015 | **A/Switzerland/9715293/2013**: E4/E2, clade 3C.3a (SNH) | 14 *(2015)* | 40; 11 | 97·76 | **L173S:1** (599), **S154A:13**  (14,306) |  | **G158K:2**(4,110), **G158R:4**(6,837),  **V202G:14**(17,342),  **I156I^+^:8**(16,900),  **I156K^+^:6**(77)  **N160K:1**(2,812), **N160S:6**(12,395),  **K176T:6**(12,943), | **S107N^M+^:1**(2,099) | **D241N:7**(928),  **I230T:1** (189), **Q213R:2**(1,111), **S175F:7** (1,050),  **S175Y:6** (13,393), **K176T:6** (12,943), | **K176T^C^:6** (12,943),  **A144T^C^:6**(13,268), **N160K^R^:1**(2,812), **N160S^R^:6**(12,395), **S140N^R^:1**(79), **S140R^R^:1** (24) | **A122T:1, D307N:1, D505N:7, D69N:3, F15S:1, G65S:1, L19F:1, L19I:7, L19V:4, L443I:1, M184V:2, N187H:1, N461S:1, Q327H:6, R342K:13, S25N:1, S278N:1, S281G:1, X235S:14, V104I:1, Y9C:1** |  |
| 2016- 2017 | **A/Hong Kong/4801/2014**: E5/E2, clade **3C.2a** (SNH) | 27 *(2016- 2017)* | 67; 11 | 97·73 | **A154S:1**  (3,053) | **S214A:2**(56), **S214P:7**(721), **S161N:2**(135), **L173S:1**(561) | **H172Q:1**(20),  **R158K:5**(4,107),  **I156K^+^:9**(73),  **S160N:13**(1,585), **N138D:1**(813),  **E78G:1**(1,492),  **E78K:1**(556),  **R158G:12**(6,183), **S160K:2**(2,812), **K176T:14**(12,910) | **S107N^M+^:1**(2,099), **N138D^M^:1**(813) | **K99E^+^:1** (129), **A212T:2**(1), **D241N:7**(667), **I230T:3** (187), **Q213R:5**(1,111), **Y175F:7** (774),  **Y175S:6** (2,916), **T144A^+^:11** (3,861),  **K176T:14**(12,910) | **T56K^R^:1*, S160N^C^:13**(1,585), **N138D^R^:1**(813),  **S140N^R^:1**(76), **T144A^R^:11** (3,861), **T151K^R^:2**(1,734) | **A492T:1, C12R:1, D509E:1, D69E:2, D69N:7, G291S:1, G495E:2, G500E:1, G94D:2, H327Q:18, I19L:10, I19V:3, I239V:2, I41V:1, I422V:7, I50V:1, I64T:2, K108R:5, K223Q:1, K275R:1, P210L:27, K292R:1, K294N:2, K342R:6, K519R:1, N137K:9, N187K:7, N461S:5, R49Q:2, N505D:12, V14G:2, Q91H:2, R421G:1, S162G:1, S112N:26, S278N:1, S328N:8, V363M:1, Y110H:2** | **I538M:2** |
| 2018 | **A/Singapore/INFIMH-16-0019/2016**: C1S3/S4, Clade **3C.2a1** (SNH) | 13 *(2017)* | 50; 13 | 96·98 | **A154S:5**  (2, 275) | **S214A:2**(33), **S214P:5**(237),  **S161N:2**(49), | **G158K:1**(3,675), **G158R:7**(3,133), **H172Q:1**(9),  **S160N:7**(119), **N138D:2**(340), **T176K:7** (2,330), **I156K^+^:5**(31), | **N138D^M^:2**(340) | **A212T:2**(1), **D241N:2**(18), **I230T:2** (149), **Q213R:1**(719), **Y175F:2** (44), **K99E^+^:1** (116), **T176K:7** (2,330), **Y175S:5** (2,168), | **S160N^C^:7**(119),  **N138D^R^: 2**(340),  **T144A^R^:5** (23), **T151K^R^:2**(1,705)  **T56K^R^:1*,** | **A492T:1, D69E:1, D69N:2, E495G:11, E500G:8, G291S:1, G94D:2, H327Q:10, I19L:7, I239V:2, I41V:1, I64T:2, K108R:3,** **K137N:8, K187N:8, K275R:1,** **K292R:2, K294N:2, K342R:5,** **N461S:1, N505D:7, Q91H:1, R421G:1, R49Q:2, S162G:1, S328N:7, V422I:8, Y110H:2, N187K:5; T144A:5** (23), | **I538M:2** |
| 2019 | **A/Switzerland/8060/2017**: E5/E1, Clade 3C.2a2 (SH) | 13 *(2017)* | 54; 16 | 97·04 | **A154S:5**  (1, 938) | **S214A:2**(22), **S214P:5**(76), **S161N:2**(25), | **H172Q:1**(3), **K147T:13 (**2,515),  **K158R:7 (**239), **I156K^+^:5**(27), **S160N:7**(11), **N138D:2**(47)  **K158G:5** (3,050), **K176T:6**(3,244) | **N138D^M^:2**(47) | **A212T:2***, **D241N:2**(1), **I230T:2** (95), **Q213R:1**(429), **Y175F:2** (44),  **K99E^+^:1** (113), **Y175S:5**(1846), **K176T:6**(3,244) | **T144A^R^:5** (2,124), **T151K^R^:2**(520)  **T56K^R^:1*** | **A492T:1, D69E:1, D69N:2,**  **G291S:1, G495E:2,**  **G500E:5, G94D:2, H327Q:10, I19L:7, I239V:2, I41V:1, I422V:5, I64T:2, K108R:3, K275R:1, K292R:1 K294N:2, K342R:5, N137K:5, N187K:5, N461S:1, N505D:7, P210L:13,** **Q277R:13, Q91H:1, R421G:1, R49Q:2, S112N:13, S162G:1, S328N:7, Y110H:2** | **I538M:2** |
|  | **A/Kansas/14/2017**: E5, Clade 3C.3a (NH) |  | 57; 18 | 96·55 | **S154A:8**  (3,435) | **S214A:2**(22), **S214P:5**(76), **S209F:13**(3,398), **S161N:2**(25) | **G158K:1**(2,052), **G158R:7**(251), **H172Q:1**(3), **I156K^+^:5**(27), **S160N:7**(11), **N138D:2**(50), **K160S**:6(3,409), **K176T:6**(3,404) | **N138D^M^:2**(50), **N107S^M+^:13**(3,513) | **N206D:13**(5,378),  **A212T:2***, **D241N:2**(1), **I230T:2** (96), **Q213R:1**(432), **S175F:2** (44), **S175Y:6** (3,505), **K176T:6**(3,404), **A144T^+^:8**(3,250), **K99E^+^:1** (113) | **N138D^R^:2**(50), **T151K^R^:2**(541), **T262N^C^:13**(5,382), **T56K^R^:1*** | **A492T:1, C9Y:13, D69E:1, D69N:2, G291S:1, G495E:2, G500E:5, G94D:2, I19L:7, I239V:2, I41V:1, I422V:5, I64T:2, K108R:3, K275R:1, K292R:1, K294N:2, M494I:13, N137K:5, N187K:5, N461S:1, N505D:7, Q327H:3, Q91H:1, R342K:8, R421G:1, R49Q:2, S162G:1, S328N:7, Y110H:2** | **I538M:2** |
| 2020 | **A/South Australia/34/2019**:  E5, Clade 3C.2a1b (SH) | 13 *(2017)* | 56; 18 | 96·36 | **A154S:5**  (1,252) | **S214A:2***, **S214P:5**(89), **S161N:2**(4) | **G158K:1**(2), **G158R:7**(13), **H172Q:1**(2),  **I202G:13**(1,954), **K147T:13 (**1,312), **I156K^+^:5**(1), **S160N:7**(59), **N138D:2**(3), **K176T:6**(3,404),  **G78E:13**(930), | **N138D^M^:2**(3), | **A212T:2***, **D241N:2***, **I230T:2** (9), **Q213R:1**(416), **Y175F:2*, K99E^+^:1** (122), **Y175S:5**(1,074), | **S160N^C^:7**(59),  **N138D^R^:2**(3), **T144A^R^:5** (1,288), **T151K^R^:2**(252), **T56K^R^:1*** | **A492T:1, D69E:1, D69N:2, E500G:8 F235S:13, G291S:1, G495E:2, G78E:13, G94D:2, I19L:7, I239V:2, I41V:1, I64T:2, K137N:8, K187N:8, K275R:1, K292R:1, K294N:2, K342R:5, M363V:13, N461S:1, N505D:7 Q327H:3, Q91H:1, R108K:10, R421G:1, R49Q:2, S162G:1, S328N:7, V422I:8, Y110H:2** | **I538M:2, I545V:13** |
| Neuraminidase (N2) | | | | | | | | | | |  |  |
| 2010-2011 | **A/Perth/16/2009**: MDCKX, A/Perth/16 clade (SNH) | 25 *(2010 - 2011)* | 32; 6·8 | 98·61 |  |  | **E221D:1**(12,894), **K369T:23**(17,155), **S367N:23** (17,347),  **E221K:1**(54), | **D251N^M^:1**(16)**,** |  | **N402D^R^:24** (15,970), **S331R^R^:11** (592) | **D127T:1, D339A:2, D93G:1, E343D:1, E344K:1, G414D:1, I26T:3, I26V:1, I307M:1, I464L:23 I77K:1, K249E:1, K308E:1, K369T:40 L338S:1, Q273R:1, R60K:3, S334N:10, T95A:3, V143M:3** | **F42L:1, L81P:16, M51V:1, N43H:1, N43Y:6, P55T:1, S44F:1, V317M:2, Y40C:2** |
| 2012 | **A/Perth/16/2009**:  MDCKX, A/Perth/16 clade (SH) | 17 *(2012)* | 19; 7 | 98·336 |  |  | **E221K:1**(54), **E344K:1**(2,739), **K369T:17**(17,155) |  |  | **N329D^R^:1**(35), **S367N^C^:40**(17,347), **N402D^R^:8** (15,970), **S331R^R^:9**(592) | **E344K:1, K369T:17,**  **D93G:5, E343D:1, I26T:9, I307M:7, I464L:17, I73V:2, I77T:1, S315N:1, S334N:9** | **L81P:8, N43H:9, V13I:2** |
|  | **A/Victoria/361/2011**:  E3/E4, Clade 3C.1 (NH) |  | 18; 6·8 | 98·11 |  |  | **T329N:5**(9,543), **E221K:1**(45), **T329D:1**(28), **E344K:1**(2,738), |  |  | **T329N^C^:5**(9,543), **D402N^C^:9** (317), | **G93D:12, E343D:1, I26T:9, I307M:7, I73V:2, I77T:1, K258E:17, S315N:1, S331R:9, S334N:9,** | **N43H:9, P81L:9, V13I:2** |
| 2013 | **A/Victoria/361/2011**:  E3/E4, Clade 3C.1 (SH) | 7 *(2013)* | 9; 2·9 | 99·25 |  |  | **T329N:5**(9,543), **T329S:2**(4,291) | **D251V^M^:1**(311)**, Y155F^S^:1**(296) |  | **T329N^C^:5**(9,543) | **G111A:1*, K258E:7, S315G:1, V165I:1** | **M241L:1** |
|  | **A/Texas/50/2012**:  E5, Clade 3C.1 (NH) |  | 8; 2 | 99·29 |  |  | **H150R:7**(14,990), **N329S:1**(4,286) | **D251V^M^:1**(297)**,**  **Y155F^S^:1**(296), **N329S^S^:1**(4,286) |  | **N329S^R^:1**(4,286) | **G111A:1*, S315G:1, V165I:1** | **M241L:1** |
| 2014 | **A/Texas/50/2012**:  E5, Clade 3C.1 (SNH) | 11 (*2014)* | 12; 2 | 99·39 |  |  | **E221D^+^:1**(12,843), **H150R:11**(14,990), **E344K:1**(2,738) | **H150R^S+^:11**(14,990), **I106V^M+^:1**(16) |  |  | **D127E:3, I392T:1, P126S:1, V143L:1** | **A82T:1, I194V:1, V317M:1, Y40C:1** |
| 2015 | **A/Switzerland/9715293/2013**: E4/E2, clade 3C.3a (SNH) | 14 *(2015)* | 19; 3·9 | 99·07 |  |  | **D221E:6**(292), **E344K:5**(2,736) | **D251V^M^:1**(195), **Y155F^S^:1**(197), **S247T ^M+^:4** (10,132), **S245N^C+^:2**(10,271) |  | **S245N^C^:2**(10,27) | **E344K:5, I28L:1, I380V:7, P468H:1, S315G:1, S315N:1, S384F:1, T267K:7, T392I:13, V231I:1, V263I:1** | **I194V:1, I30V:1, M51T:2** |
| 2016- 2017 | **A/Hong Kong/4801/2014**: E5/E3, clade **3C.2a** (SNH) | 27 *(2016- 2017)* | 49; 7·8 | 98·10 |  |  | **D221E:7**(215), **N329S:1**(4,286),  **E344K:3**(2,735), **G346V:1**(289), **K220N:1**(1,604), **R400K:1**(80), **K220R:1**(2), **R150H:1**(178) | **D251V^M^:1**(133), **Y155F^S^:1**(133), **S247T ^M+^:10** (10,130), **R150H^S+^:1**(178) |  | **N329S^R^:3**(4,283),  **D309N^C^:1**(71), **D402N^C^:2** (33), **N86H^R^:1**(1), **S245N^C+^:2**(10,268), **S331R^R^:6**(527) | **D339N:6, G286D:1, G93D:4, H347N:1**, **I176M:2, I231V:27, I26V:6, I307M:2, I380V:13, I65V:1, K249E:6, K431R:1, L338S:6, L464I:1, P126L:1, P468H:7, P468L:2, Q273R:6, S315G:1, S315N:1, T267K:14, T392I:27, T69A:1, T95A:7** | **F23L:2, I30V:1, M51V:6, N161S:2, N43Y:2,** **P55T:1, P81L:2,** **P81S:1, V303I:2** |
| 2018 | **A/Singapore/INFIMH-16-0019/2016**: C1S3/S4, Clade **3C.2a1** (SNH) | 13 *(2017)* | 30; 9 | 97·28 |  |  | **D221E^+^:2**(4), **N329S:2**(4,267),  **K220R:1**(2), | **N245S^M+^:7**(257), **T247S ^M+^:7**(240), |  | **N329S^R^:2**(4,267), **N245S^R^:7**(257), **D402N^C^:2** (16), **N86H^R^:1**(1), **S331R^R^:5**(135) | **G93D:2, H347N:1**, **H468P:7, I176M:2, I26V:5, I212V:13,**  **I307M:1, K249E:5, K267T:7, L338S:5, L464I:1, N339D:10, Q273R:5, T95A:6, V380I:7** | **I30V:1, M51V:5, N43Y:2, P55T:1, P81L:2, P81S:1,** **V303I:1** |
| 2019 | **A/Switzerland/8060/2017**: E5/E2, Clade 3C.2a2 (SH) | 13 *(2017)* | 31; 11 | 96·78 |  |  | **D221E^+^:2***, **S329N:11**(81),  **K220R:1**(2), | **N245S^M+^:7**(28), **T247S ^M+^:7**(11) |  | **S329N^C^:11**(81), **N245S^R^:7**(28), **D402N^C^:2** (1), **N86H^R^:1**(1) | **G93D:2, H347N:1**, **H468P:7, I26V:1, I307M:1, K249E:5, K267T:7, L338S:5, L464I:1, M176I:11, N339D:10, Q273R:5, S331R:5,** **S386P:13, T95A:6, V380I:7** | **I30V:1, M51V:5, N43Y:2, P55T:1, P81L:2, P81S:1,** **V194I:13, V303I:1** |
|  | **A/Kansas/14/2017**: E5, Clade 3C.3a (NH) |  | 35; 13·8 | 96·49 |  |  | **D221E^+^:2***, **T329N:11**(89),  **K220R:1**(2), **K344E:13**(1,819), **T329S:2**(2,519) | **T247S^M+^:7**(11), **N245S^M+^:7**(29), **A149V^M^:13**(2,578), **H155Y^S^:13**(2,598), |  | **T329N^C^:11**(89), **N245S^R^:7**(29), **D402N^C^:2** (1), **N86H^R^:1**(1) | **G93D:2, H347N:1**, **H468P:7, I140L:13, I176M:2, I26V:5, I307M:1, K249E:5, K267T:7, L338S:5, L464I:1, N339D:10, Q273R:5, R75K:13, S331R:5, T95A:6, V380I:7** | **I30V:1, M51V:5, N43Y:2, P55T:1, P81L:2, P81S:1, V303I:1** |
| 2020 | **A/South Australia/34/2019**:  E5, Clade 3C.2a1b (SH) | 13 *(2017)* | 33; 13·5 | 96·55 |  |  | **D221E^+^:2***,  **S329N:11**(9),  **N220K:12**(1,116), **N220R:1**(1),  **K344E:13**(84), | **T247S^M+^:7***, **N245S^M+^:7**(5) |  | **S329N^C^:11**(9), **N245S^R^:7**(5), **D402N^C^:2** (3), **N86H^R^:1*** | **G93D:2, H347N:1**, **H468P:7, I176M:2, I26V:5, I307M:1, K249E:5, K267T:7, L126P:13, L338S:5, L464I:1, N339D:10, Q273R:5, R315S:13, S331R:5, T95A:6, V380I:7** | **I303V:12, I30V:1, M51V:5, N43Y:2, P55T:1, P81L:2, P81S:1** |
| Matrix Protein (M1 and M2) | | | | | | | | | | |  |  |
| 2010-2011 | **A/Perth/16/2009**: MDCKX, A/Perth/16 clade (SNH) | 25 *(2010 - 2011)* | 6: 1 | 99·86 |  |  |  |  | **G16E:1**(759) |  | **P25Q:1, M93I:1, K98R:1, L40F:1*** | **D88E:1, R18K:2** |
| 2012 | **A/Perth/16/2009**: MDCKX, A/Perth/16 clade (SH) | 17 *(2012)* | 4; 1 | 99·55 |  |  |  |  |  |  | **F48S:1**, **K98R:1, R61G:3** | **D88E:9** |
|  | **A/Victoria/361/2011**:  E3/E4, Clade 3C.1 (NH) |  | 4; 1 | 99·55 |  |  |  |  |  |  | **F48S:1**, **K98R:1, R61G:3** | **D88E:9** |
| 2013 | **A/Victoria/361/2011**:  E3/E4, Clade 3C.1 (SH) | 7 *(2013)* | 4; 1 | 99·78 |  |  |  |  |  |  | **A33V:1, F48S:1**, **L54V:1** | **V15I:1** |
|  | **A/Texas/50/2012**:  E5, Clade 3C.1 (NH) |  | 4; 1 | 99·78 |  |  |  |  |  |  | **A33V:1, F48S:1**, **L54V:1** | **V15I:1** |
| 2014 | **A/Texas/50/2012**:  E5/E1, Clade 3C.1 (SNH) | 12 (*2014)* | 4; 1 | 99·33 |  |  |  |  |  |  | **F48S:7, I39M:1** | **E14G:1, T227A:1** |
| 2015 | **A/Switzerland/9715293/2013**: E4/E2, clade 3C.3a (SNH) | 14 *(2015)* | 7; 1 | 99·14 |  |  |  | **V27A^SM+^:2**(22), |  |  | **F48S:7, I39M:3, L54V:1** | **N20S:1, Q208R:1, T239A:2** |
| 2016- 2017 | **A/Hong Kong/4801/2014**: E5/E3, clade **3C.2a** (SNH) | 29 *(2016- 2017)* | 15; 1 | 99·08 |  |  |  | **V27A^SM+^:6**(21), | **G16E:7**(483) |  | **D21G:1, F48S:10, L54V:1, L59I:1, R61K:1** | **E235K:1, K242R:1, N13D:1, N20S:1, R18K:1, T227A:1, T227N:1, T239A:6** |
| 2018 | **A/Singapore/INFIMH-16-0019/2016**: C1S3/S4, Clade **3C.2a1** (SNH) | 14 *(2017)* | 8; 1 | 98·49 |  |  |  | **A27V^SM+^:13**(6,981) | **G16E:6**(132) |  | **D21G:1, R61K:1, F48S:1** | **N20S:1, R18K:1, T239A:1** |
| 2019 | **A/Switzerland/8060/2017**: E5/E2, Clade 3C.2a2 (SH) | 14 *(2017)* | 8; 1 | 99·28 |  |  |  | **V27A^SM+^:1**(1), **G16E ^S+^:6**(24) | **G16E:6**(24) |  | **D21G:1, R61K:1, F48S:1** | **N20S:1, R18K:1, T239A:1** |
|  | **A/Kansas/14/2017**: E5, Clade 3C.3a (NH) |  | 10; 1·89 | 97·48 |  |  |  | **I27A:1**(1),  **I27V:13**(2,635), **G16E^S+^:6**(24) | **G16E:6**(24) |  | **C52Y:14, D21G:1, F48S:1, R61K:1** | **N20S:1, R18K:1, T239A:1** |
| 2020 | **A/South Australia/34/2019**:  E5, Clade 3C.2a1b (SH) | 14 *(2017)* | 9; 1 | 98·34 |  |  |  | **V27A^SM+^:1***, **G16E^S+^:6**(15) | **G16E:6**(15) |  | **D21G:1, F48S:1, R61K:1, L25P:14,** | **N20S:1, R18K:1, T239A:1** |
| Data are in n or n (%), unless otherwise indicated. AA= Amino acid. AAS= Amino acid substitutions. SH= Southern hemisphere vaccine. HN= Northern hemisphere vaccine. SNH= vaccine strain shared by both the Southern and Northern hemispheres for a given influenza season. The number of unique amino acid substitutions observed in all proteins sequences and per protein sequence are represented as N; n. The number of times a substitution is observed in the Uganda virus proteins and globally is reported in BOLD and (bracket), respectively. Substitutions are colour-coded based on their known or predicted biological function and level of significance. Red mutations are the most significant (interestlevel =3) as they alter virulence, cause strong drug resistance and reverse premature stop codon in PB1-F2. The Orange (significant, interestlevel=2) occur at drug binding sites, affect host specificity and cause antigenic shift and mild drug resistance. Magenta (significant, interestlevel=2) adds or removes glycosylation sites. Blue mutations (moderately significant, interestlevel=1) have structural functions at interaction sites. Structural functions include host cell receptor binding, binding small ligand(s), viral oligomerization interfaces, and antibody recognition sites. The Green mutations (least significant, interestlevel=0) are common to subtypes while Black (least significant with warnlevel=0) have no known effects. Superscript symbol definitions: “*” = AAS reported for the first time (current global frequency = 0). “^+^” = AAS have their function reported in combination with others. “^R^” = AAS removes a potential glycosylation site. “^C^” = AAS creates a potential glycosylation site. “^S^” = AAS causes strong drug resistance. “^M^” = AAS causes mild drug resistance. “^@^” = AAS is a common subtype marker. | | | | | | | | | | | | |

**sTable 5: Amino acid substitutions in the complete coding sequences of HA (H3), NA (N2) and MP (M1 and M2) of Uganda A(H3N2) viruses compared to Southern and Northern Hemisphere vaccines**. Amino acid similarity, amino acid substitutions and their corresponding functions and frequencies were obtained from Flusurver (<http://flusurver.bii.a-star.edu.sg>; accessed on 30^th^ March 2021).

**sFigure 5: Multiple sequence alignment (MSA) of Uganda A(H3N2) M2 genes with adamantine-susceptible A/New York/392/2004(H3N2)**

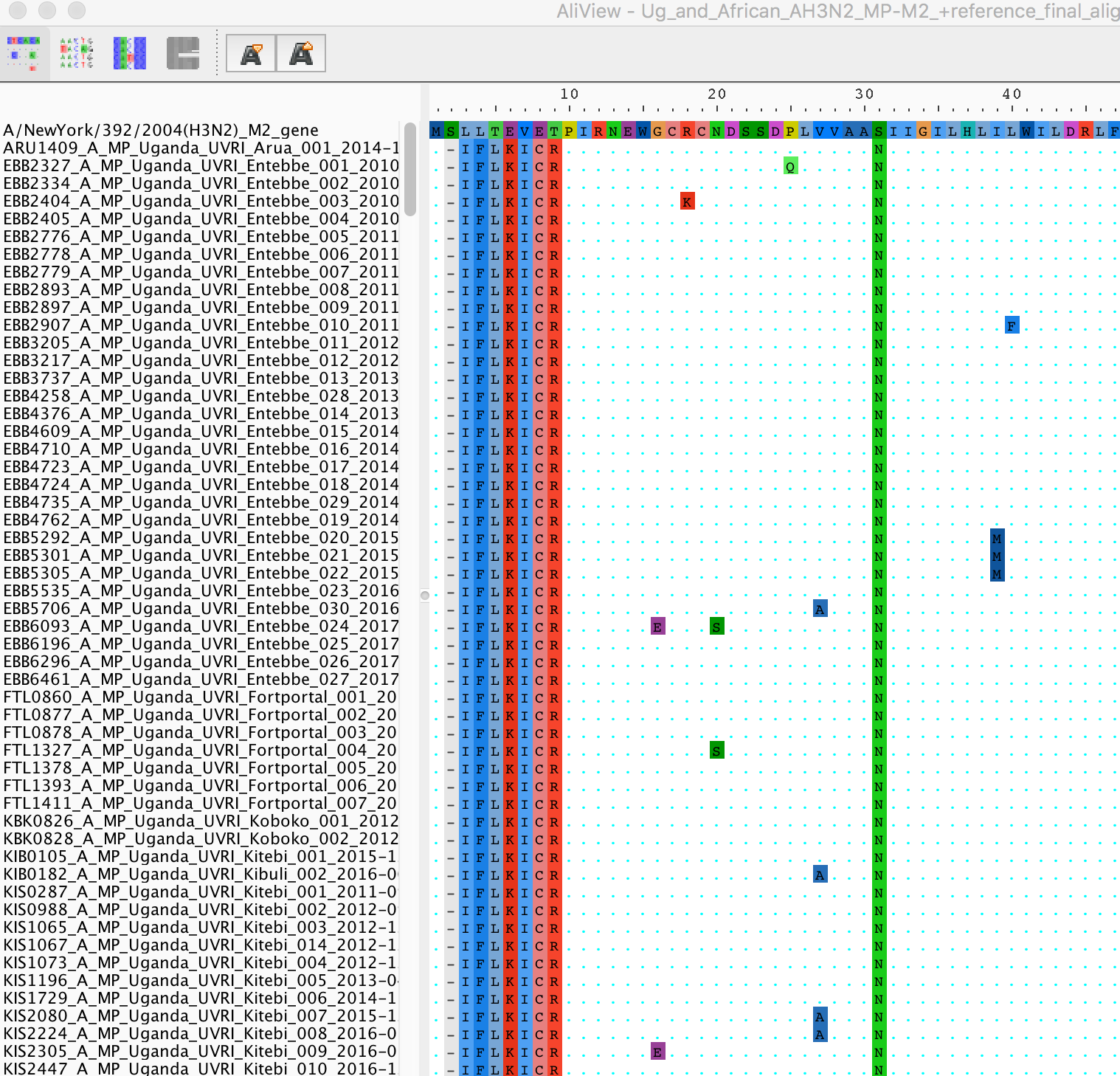

**SFigure 5: Multiple sequence alignment of Uganda A(H3N2) M2 genes with adamantine-susceptible A/New York/392/2004(H3N2) as a reference.** Substitutions with Alanine (A) and Asparagine (N) are observed in Uganda viruses at positions 27 and 31, respectively. Mutations were visualized using AliView (Larsson, 2014)^8^.

#### **sFigure 6: Phylogenies showing the spatial divergence Uganda influenza A viruses**

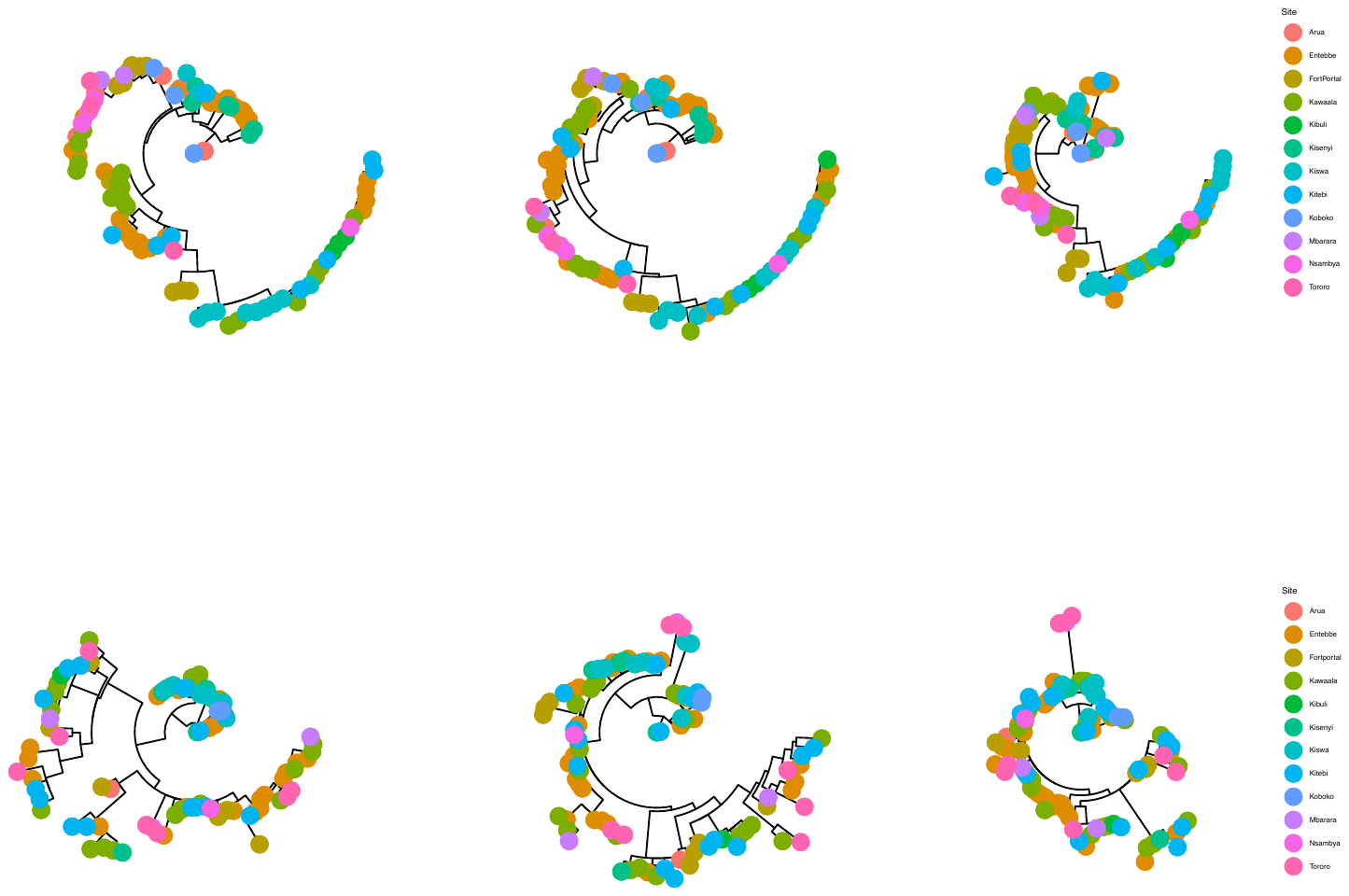

**sFigure 6: Phylogenies showing the spatial divergence of the HA, NA and MP genes of Uganda A(H1N1)pdm09 (A) and A(H3N2) (B) influenza viruses collected from 2010 to 2018**. Trees were rooted using the oldest sequence in the Ugandan dataset. There was no uniqueness in viruses circulating in a given site or geographical region in all gene trees.

#### **sFigure 7: Phylogenetic clustering of Uganda and other Africa A(H1N1)pdm09 viruses in the hemagglutinin gene**

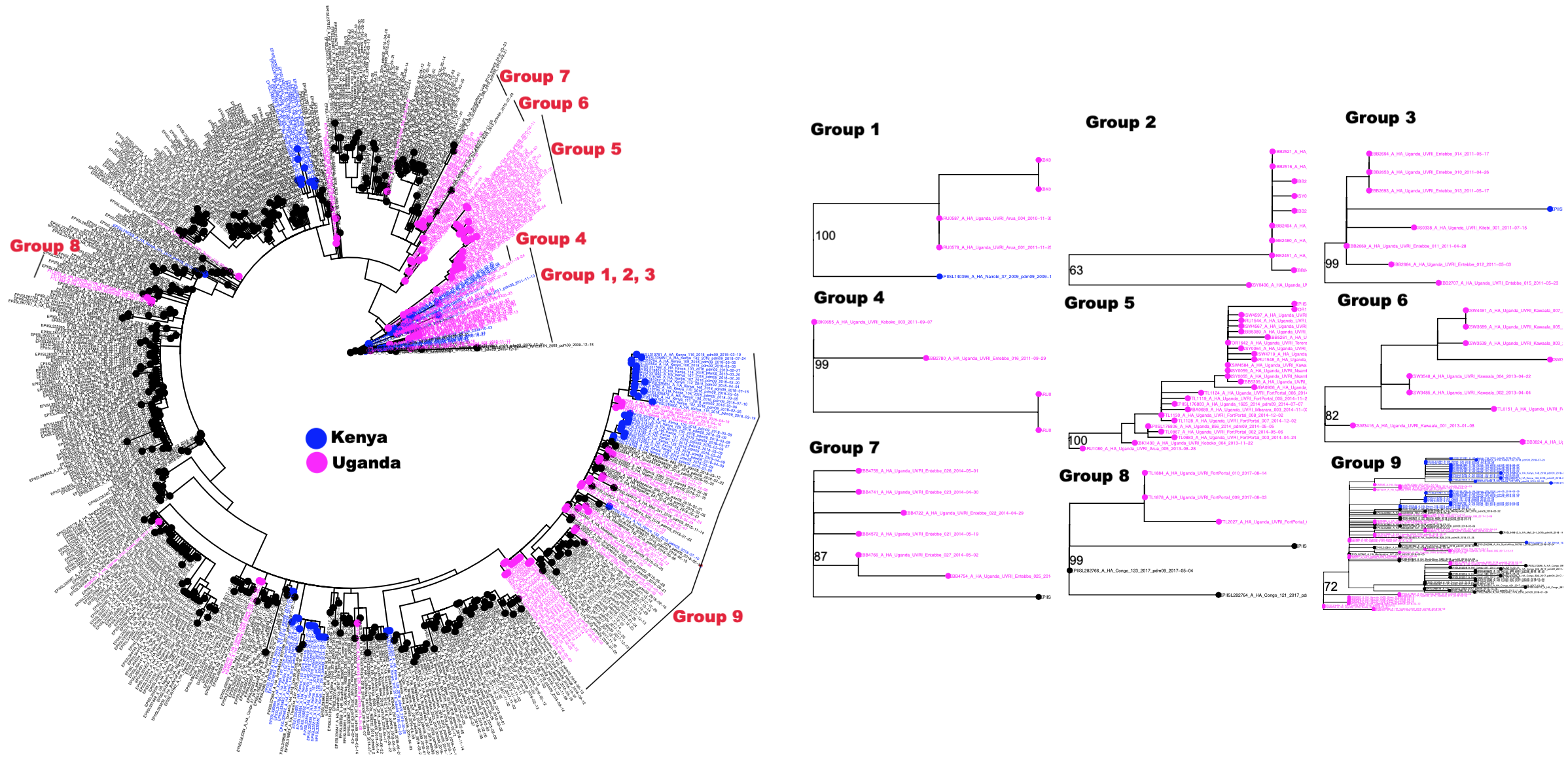

**sFigure 7:** **Phylogenetic clustering of 2009-2018 Uganda A(H1N1)pdm09 viruses with other African viruses (2009-2019) in the HA (H1) gene.** Groups are defined as phylogenetic clusters with 3 or more Uganda virus sequences. All 9 groups are shown. Group details support are described in sTable 6A below.

#### **sFigure 8: Phylogenetic clustering of Uganda and other Africa A(H1N1)pdm09 viruses in the neuraminidase gene**

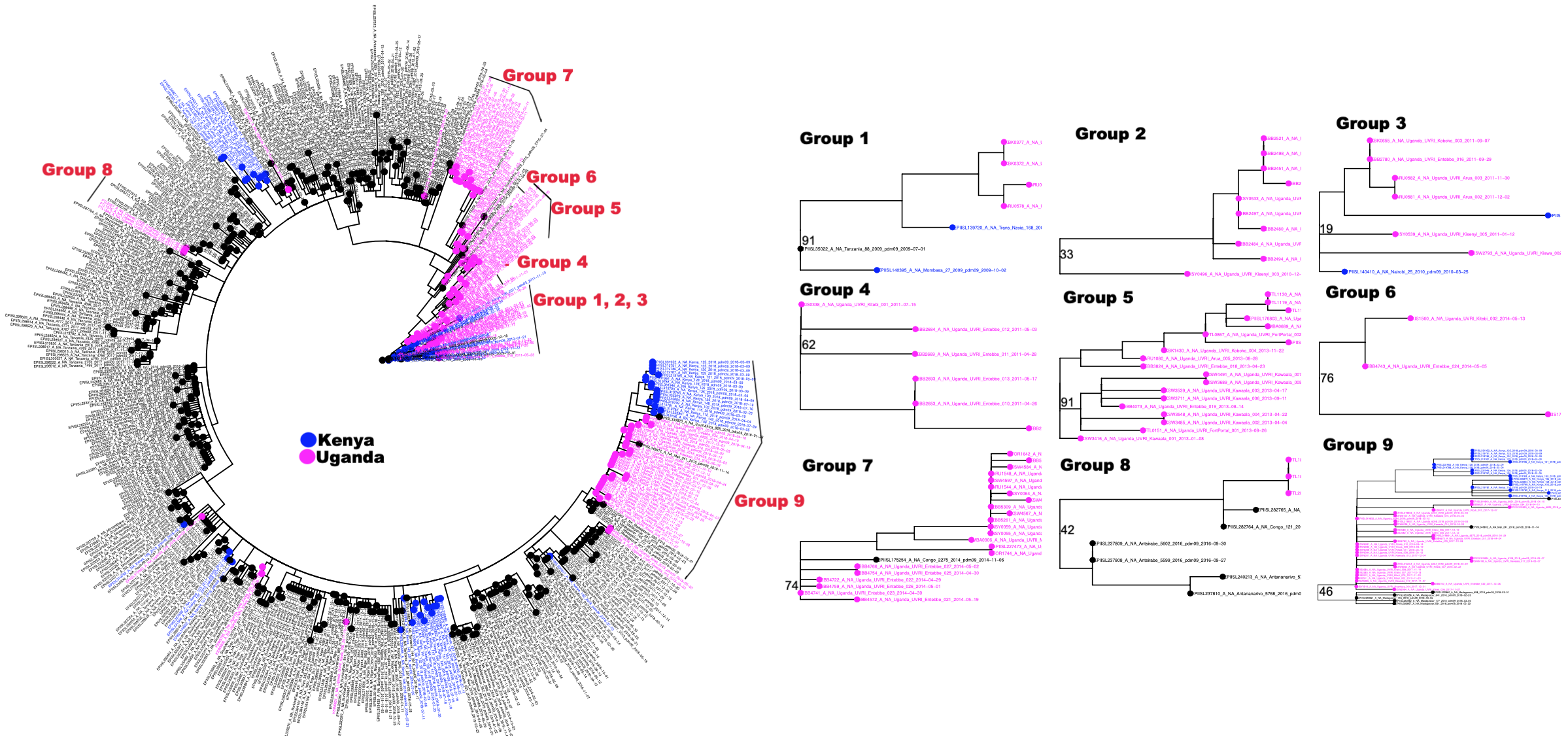

**sFigure 8: Phylogenetic clustering of 2009-2018 Uganda A(H1N1)pdm09 viruses with other African viruses (2009-2019) in the NA (N1) gene.** All 9 groups are shown. Group details support are described in sTable 6A below.

#### **sFigure 9: Phylogenetic clustering of Uganda and other Africa A(H1N1)pdm09 viruses in the matrix protein gene**

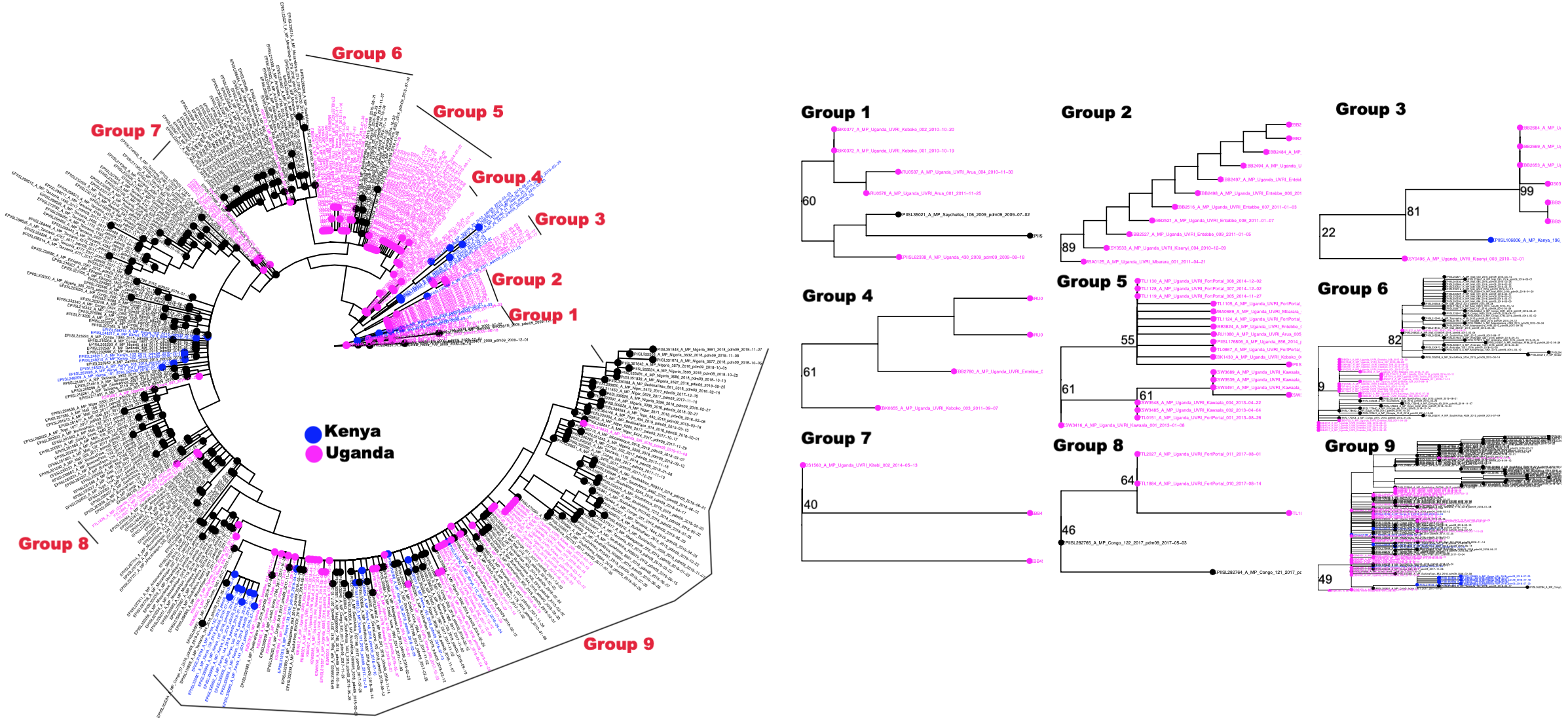

**sFigure 9:** **Phylogenetic clustering of 2009-2018 Uganda A(H1N1)pdm09 viruses with other African viruses (2009-2019) in the MP gene.** All 9 groups are shown. Group details support are described in sTable 6A below.

#### **sFigure 10: Phylogenetic clustering of Uganda and other Africa A(H3N2) viruses in the hemagglutinin gene**

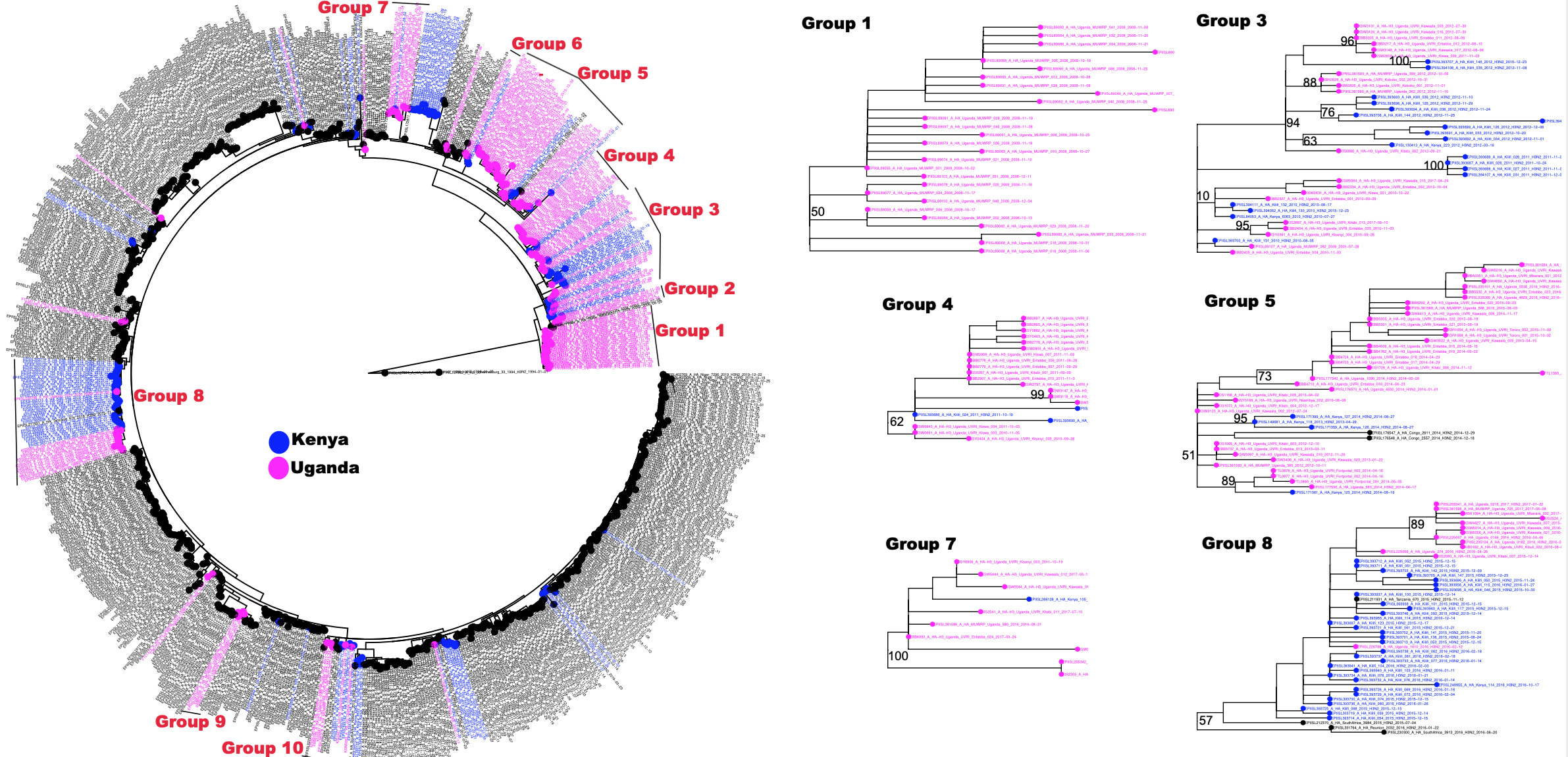

**sFigure 10: Phylogenetic clustering of 2008- 2017 Uganda A(H3N2) viruses with other African viruses (1994-2019) in the HA (H3) gene.** There were 10 groups. Only the large 6 groups are shown. The MWRP 2008 and 2009 viruses dominated H3 groups 1 and 2, respectively. Group 2 (not shown) contained only 3 Uganda viruses isolated in 2009 by MWRP. Group details support are described in sTable 6B below.

#### **sFigure 11:** **Phylogenetic clustering of Uganda and other Africa A(H3N2) viruses in the neuraminidase gene**

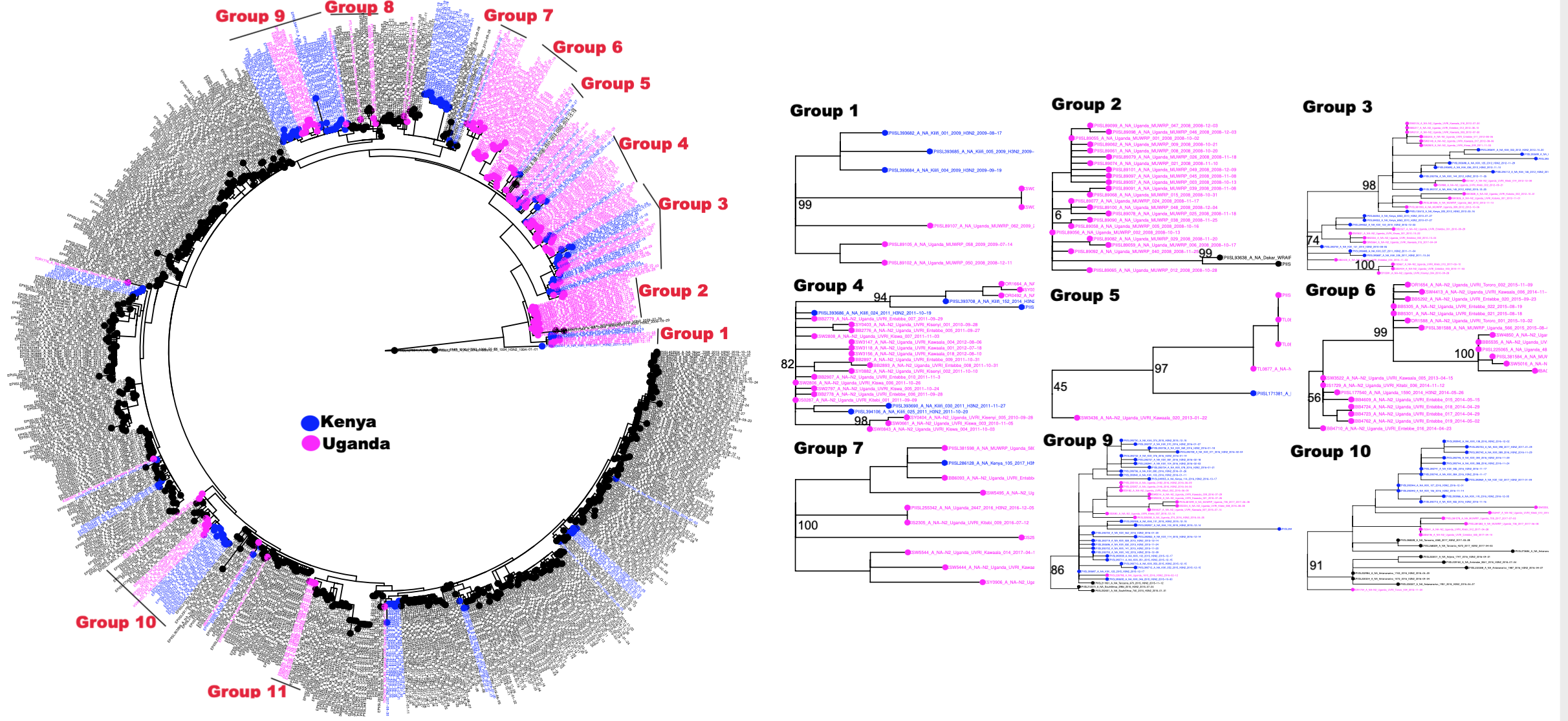

**sFigure 11:** **Phylogenetic clustering of 2008-2017 Uganda A(H3N2) viruses with other African viruses (1994-2019) in the NA (N2) gene.** Only the large groups (9/11) with 5 or more Uganda sequences are shown. Group details support are described in sTable 6B below.

#### **sFigure 12:** **Phylogenetic clustering of Uganda and other Africa A(H3N2) viruses in the matrix protein gene**

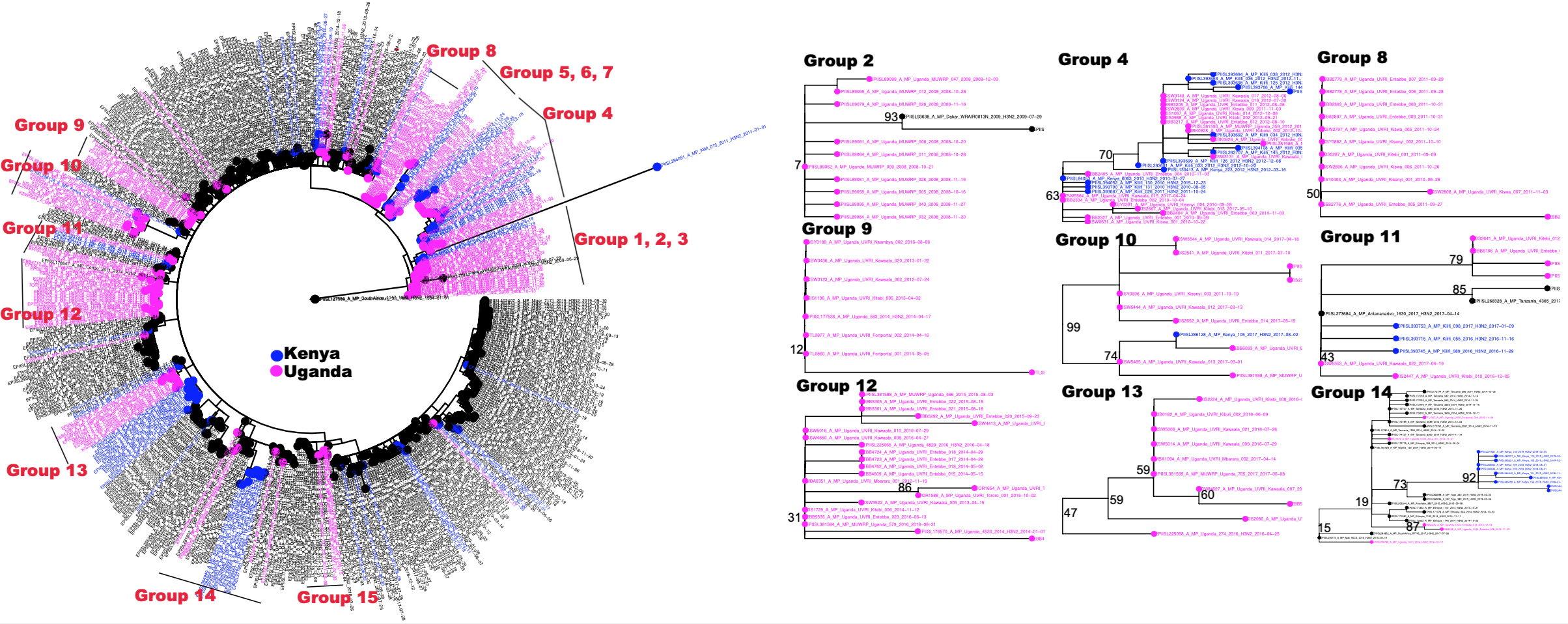

**sFigure 12:** **Phylogenetic clustering of 2008-2017 Uganda A(H3N2) viruses with other African viruses (1994-2019) in the MP gene.** Only the large groups (9/15) with 5 or more Uganda sequences are displayed. Group details support are described in sTable 6B below.

#### **sTable 6: Details of phylogenetic groups among the newly-generated Uganda viruses and other African viruses**

| Group | Bootstrap | Number of Uganda viruses/ total number of viruses in the group | Origin of other  viruses | Years sampled | Group | Bootstrap | Number of Uganda viruses/ total number of viruses in the group | Origin of other viruses | Year sampled | Group | Bootstrap | Number of Uganda viruses/ total number of viruses in the group | Origin of other viruses | Year sampled |
| --- | --- | --- | --- | --- | --- | --- | --- | --- | --- | --- | --- | --- | --- | --- |
| A. Hemagglutinin (H1) | | | | | **Neuraminidase (N1)** | | | | | **Matrix protein (MP)** | | | | |
| 1 | 100 | 4/5 | Nairobi | 2009 -2011 | **1** | 91 | 4/7 | 2010 | 2009 - 2011 | **1** | 60 | 5/7 | Bamako, Seychelles | 2009 – 2011 |
| 2 | 63 | 10/10 | - | 2010 -2011 | **2** | 33 | 10/10 | - | 2010 - 2011 | **2** | 89 | 11/11 | - | 2010 -2011 |
| 3 | 99 | 7/8 | Kenya | 2011 | **3** | 19 | 6/8 | Nairobi, Kenya | 2010 - 2011 | **3** | 22 | 7/8 | Kenya | 2010 - 2011 |
| 4 | 99 | 4/4 | - | 2011 | **4** | 62 | 6/6 | - | 2011 | **4** | 61 | 4/4 | - | 2011 |
| 5 | 100 | 27/27 | - | 2013 -2016 | **5** | 91 | 19/19 | - | 2013 - 2015 | **5** | 61 | 20/20 | - | 2013 - 2015 |
| 6 | 82 | 9/9 | - | 2013, 2015 | **6** | 76 | 3/3 | - | 2014 | **6** | 9 | 22/64 | Congo, South Africa, Ethiopia, Egypt, Burkina Faso, Mozambique, Moramanga, Antsirabe, Mali, Maevatanana, NosyBe, Toamasina | 2014 - 2016 |
| 7 | 87 | 6/7 | Congo | 2014 | **7** | 74 | 22/23 | Congo | 2014 -2016 | **7** | 40 | 3/3 | - | 2014 |
| 8 | 99 | 3/6 | Congo | 2017 | **8** | 42 | 3/9 | Antsirabe (Madagascar), Antananarivo (Madagascar), Congo, | 2016 - 2017 | **8** | 46 | 3/5 | Congo | 2017 |
| 9 | 72 | 32/ 93 | Kenya, Madagascar, South Africa, Nigeria, Mali, Congo, Tanzania | 2017-2018 | **9** | 46 | 31/55 | Madagascar, Mali, South Africa, Kenya | 2017 -2018 | **9** | 49 | 34/128 | CoteD’iviore, Congo, Tanzania, Kenya, South Africa, Madagascar, Mali, Burkina Faso, Sierra leone, Togo, Nigeria, Mozambique, Niger | 2017 -2019 |
| B. Hemagglutinin (H3) | | | | | **Neuraminidase (N2)** | | | | | **Matrix protein (MP)** | | | | |
| 1 | 50 | 28/28 | - | 2008 | **1** | 99 | 5/8 | Kilifi | 2008 - 2010 | **1** | 56 | 3/3 | - | 2008 |
| 2 | 92 | 3/3 | - | 2009 | **2** | <10 | 22/24 | Dakar | 2008 - 2009 | **2** | <10 | 10/12 | Dakar | 2008 - 2009 |
| 3 | 10 | 20/39 | Kilifi, Kenya | 2009 – 2012, 2015, 2017 | **3** | 74 | 20/35 | Kilifi, Kenya | 2010 - 2012, 2015 | **3** | 14 | 5/5 | - | 2008 - 2009 |
| 4 | 62 | 18/21 | Kilifi | 2010 -2012 | **4** | 82 | 21/26 | Kilifi | 2010 - 2016 | **4** | 63 | 20/35 | Kilifi, Kenya | 2010- 2012, 2015, 2017 |
| 5 | 51 | 38/ 42 | Congo, Kenya | 2012 - 2016 | **5** | 45 | 5/6 | Kenya | 2013 - 2014 | **5** | 61 | 3/3 | - | 2010 -2011 |
| 6 | 97 | 3/5 | Kilifi, Kenya | 2013 - 2016 | **6** | 56 | 21/21 | - | 2013 - 2016 | **6** | 89 | 3/3 | - | 2012 |
| 7 | 100 | 9/10 | Kenya | 2011, 2016 -2017 | **7** | 100 | 9/10 | Kenya | 2011, 2016-2017 | **7** | 96 | 3/5 | Kilifi, Kenya | 2013 - 2016 |
| 8 | 57 | 13/48 | Reunion, South Africa, Kilifi, Kenya, Tanzania | 2015 - 2017 | **8** | 91 | 3/14 | Congo, Tanzania, Rwanda | 2014 - 2015 | **8** | 50 | 12/12 | - | 2011 |
| 9 | 96 | 4/11 | Congo | 2017 - 2018 | **9** | 86 | 11/ 40 | South Africa, Tanzania, Kilifi, | 2015 - 2017 | **9** | 12 | 8/8 | - | 2012 - 2014, 2016 |
| 10 | 51 | 4/5 | Madagascar | 2017 - 2018 | **10** | 91 | 7/28 | Anjeva, Antsirabe, Antananarivo, Kilifi, Tanzania | 2015 - 2017 | **10** | 99 | 10/11 | Kenya | 2016 - 2017 |
|  |  |  |  |  | **11** | 60 | 4/11 | Congo | 2017 | **11** | 43 | 6/12 | Antananarivo, Kilifi, Tanzania | 2016 - 2017 |
|  |  |  |  |  |  |  |  |  |  | **12** | 31 | 21/21 | - | 2014 - 2016 |
|  |  |  |  |  |  |  |  |  |  | **13** | 47 | 10/10 | - | 2016 - 2017 |
|  |  |  |  |  |  |  |  |  |  | **14** | 15 | 5/36 | Mali, South Africa, Ethiopia, Antsirabe, Togo, Kenya, Nigeria, Tanzania | 2013 - 2015, 2018 - 2019 |
|  |  |  |  |  |  |  |  |  |  | **15** | 53 | 4/10 | Congo | 2017 |

**sTable 6: Phylogenetic groups of Uganda A(H1N1)pdm09 and A(H3N2) viruses isolated in 2008-2018 [including (2008-2009) Makerere Walter Reed project (MWRP) and our new (2010-2018) dataset] with other African viruses (1994-2019) in the HA, NA, and MP genes**. The number of Uganda viruses, country or city of origin for other African viruses, and years of viral sampling are reported for each group per gene. Panel A. Panel B group 1, 2 and 3 consisted of the MWRP sequences.

### **sTable 7:** **Accession numbers vaccine strains and clade references and Africa sequences analysed in this study**

|  | A(H1N1)pdm09 | A(H3N2) |
| --- | --- | --- |
| Vaccine and clade reference viruses | EPIISL368243, EPIISL203615, EPIISL29712, EPIISL90718, EPIISL139667, EPIISL93746, EPIISL29955, EPIISL30050, EPIISL79721, EPIISL90787, EPIISL101586, EPIISL101589, EPIISL101558, EPIISL123461, EPIISL90760, EPIISL145447, EPIISL199532, EPIISL206099, EPIISL284660, EPIISL294119, EPIISL344858, EPIISL332840, EPIISL321313, EPIISL338060, EPIISL403338, EPIISL145446, EPIISL145423, EPIISL89916, EPIISL99815, EPIISL99900, EPIISL99899, EPIISL391019 | EPIISL31055, EPIISL66566, EPIISL93711, EPIISL86071, EPIISL93712, EPIISL94725, EPIISL107823, EPIISL165578, EPIISL132908, EPIISL165829, EPIISL109761, EPIISL158723, EPIISL270160, EPIISL136375, EPIISL168694, EPIISL193193, EPIISL330262, EPIISL208201, EPIISL275853, EPIISL314925, EPIISL292377, EPIISL312267, EPIISL331821, EPIISL331821, EPIISL332807, EPIISL286866, EPIISL143568, EPIISL156805, EPIISL143559, EPIISL145494, EPIISL312041, EPIISL292575, EPIISL162145, EPIISL166796, EPIISL93709, EPIISL223632, EPIISL175051, EPIISL88034, EPIISL90628, EPIISL79335, EPIISL78687, EPIISL93788, EPIISL96037, EPIISL355880, EPIISL96037 |
| Africa viruses | "EPIISL106806" "EPIISL139708" "EPIISL139716" "EPIISL139719" "EPIISL139720" "EPIISL140396" "EPIISL140404" "EPIISL140405" "EPIISL140406" "EPIISL140407" "EPIISL140408" "EPIISL140409" "EPIISL140410" "EPIISL140411" "EPIISL140412" "EPIISL140413" "EPIISL171814" "EPIISL172640" "EPIISL172641" "EPIISL172648" "EPIISL175254" "EPIISL175259" "EPIISL175880" "EPIISL176803" "EPIISL176806" "EPIISL178482" "EPIISL178484" "EPIISL203293" "EPIISL203300" "EPIISL205286" "EPIISL205287" "EPIISL205288" "EPIISL205293" "EPIISL205294" "EPIISL205297" "EPIISL205298" "EPIISL205486" "EPIISL205487" "EPIISL206563" "EPIISL206565" "EPIISL206566" "EPIISL207384" "EPIISL207387" "EPIISL207922" "EPIISL207923" "EPIISL207929" "EPIISL207930" "EPIISL207947" "EPIISL207950" "EPIISL210330" "EPIISL210332" "EPIISL210340" "EPIISL211950" "EPIISL211951" "EPIISL211952" "EPIISL213141" "EPIISL213145" "EPIISL213146" "EPIISL213147" "EPIISL213208" "EPIISL214171" "EPIISL214908" "EPIISL214909" "EPIISL214910" "EPIISL214911" "EPIISL214912" "EPIISL216223" "EPIISL216224" "EPIISL216226" "EPIISL216227" "EPIISL216256" "EPIISL216257" "EPIISL216258" "EPIISL216261" "EPIISL216262" "EPIISL216284" "EPIISL216285" "EPIISL216286" "EPIISL216287" "EPIISL218134" "EPIISL218135" "EPIISL218177" "EPIISL220958" "EPIISL220960" "EPIISL220962" "EPIISL220965" "EPIISL220967" ] "EPIISL220969" "EPIISL220999" "EPIISL221004" "EPIISL221006" "EPIISL221008" "EPIISL221010" "EPIISL221654" "EPIISL223223" "EPIISL223224" "EPIISL223226" "EPIISL223227" "EPIISL223229" "EPIISL223232" "EPIISL227473" "EPIISL230453" "EPIISL230457" "EPIISL230459" "EPIISL230461" "EPIISL230465" "EPIISL230467" "EPIISL230470" "EPIISL230471" "EPIISL230473" "EPIISL230529" "EPIISL230531" "EPIISL230532" "EPIISL230534" "EPIISL230535" "EPIISL230536" "EPIISL230537" "EPIISL232134" "EPIISL232136" "EPIISL232587" "EPIISL232588" "EPIISL232589" "EPIISL232590" "EPIISL232591" "EPIISL232592" "EPIISL232603" "EPIISL232604" "EPIISL232605" "EPIISL232606" "EPIISL232607" "EPIISL232664" "EPIISL232666" "EPIISL232667" "EPIISL232668" "EPIISL232669" "EPIISL232672" "EPIISL232674" "EPIISL232677" "EPIISL232678" "EPIISL232679" "EPIISL232680" "EPIISL232683" "EPIISL232684" "EPIISL232686" "EPIISL232688" "EPIISL233022" "EPIISL233023" "EPIISL233062" "EPIISL233063" "EPIISL233065" "EPIISL233066" "EPIISL233489" "EPIISL233490" "EPIISL235013" "EPIISL235015" "EPIISL235016" "EPIISL235017" "EPIISL235339" "EPIISL235340" "EPIISL235341" "EPIISL235342" "EPIISL235343" "EPIISL235344" "EPIISL235345" "EPIISL235346" "EPIISL236216" "EPIISL236217" "EPIISL236837" "EPIISL237808" "EPIISL237809" "EPIISL237811" "EPIISL237813" "EPIISL237822" "EPIISL240213" "EPIISL248209" "EPIISL248210" "EPIISL248211" "EPIISL248212" "EPIISL248213" "EPIISL248216" "EPIISL248217" "EPIISL256091" "EPIISL256092" "EPIISL268443" "EPIISL268444" "EPIISL268448" "EPIISL268456" "EPIISL268462" "EPIISL273782" "EPIISL275663" "EPIISL275664" "EPIISL275665" "EPIISL275666" "EPIISL275667" "EPIISL281577" "EPIISL281578" "EPIISL281579" "EPIISL281581" "EPIISL281583" "EPIISL281585" "EPIISL281587" "EPIISL281588" "EPIISL281589" "EPIISL281590" "EPIISL281613" "EPIISL281616" "EPIISL281618" "EPIISL281619" "EPIISL281620" "EPIISL281621" "EPIISL281622" "EPIISL281623" "EPIISL281624" "EPIISL281625" "EPIISL281626" "EPIISL281629" "EPIISL282764" "EPIISL282765" "EPIISL282766" "EPIISL282771" "EPIISL282776" "EPIISL283210" "EPIISL283212" "EPIISL283213" "EPIISL283214" "EPIISL283215" "EPIISL283217" "EPIISL283218" "EPIISL283219" "EPIISL283221" "EPIISL283222" "EPIISL283223" "EPIISL283225" "EPIISL283226" "EPIISL283227" "EPIISL283229" "EPIISL283231" "EPIISL283233" "EPIISL283235" "EPIISL283237" "EPIISL285866" "EPIISL287686" "EPIISL287687" "EPIISL287688" "EPIISL287704" "EPIISL287706" "EPIISL287707" "EPIISL287708" "EPIISL288566" "EPIISL289562" "EPIISL292622" "EPIISL292623" "EPIISL292624" "EPIISL292625" "EPIISL292626" "EPIISL292627" "EPIISL292628" "EPIISL292629" "EPIISL292630" "EPIISL292631" "EPIISL296234" "EPIISL296235" "EPIISL296237" "EPIISL298512" "EPIISL298513" "EPIISL298514" "EPIISL298515" "EPIISL298516" "EPIISL298517" "EPIISL298518" "EPIISL298519" "EPIISL298520" "EPIISL298522" "EPIISL298524" "EPIISL298525" "EPIISL298526" "EPIISL298528" "EPIISL298529" "EPIISL299835" "EPIISL299837" "EPIISL299838" "EPIISL299845" "EPIISL299846" "EPIISL299849" "EPIISL299850" "EPIISL299851" "EPIISL300845" "EPIISL300849" "EPIISL300851" "EPIISL300852" "EPIISL300853" "EPIISL300854" "EPIISL300855" "EPIISL300857" "EPIISL300859" "EPIISL300885" "EPIISL300901" "EPIISL300903" "EPIISL300909" "EPIISL300913" "EPIISL300916" "EPIISL305251" "EPIISL305252" "EPIISL305253" "EPIISL305254" "EPIISL305265" "EPIISL305266" "EPIISL305267" "EPIISL305269" "EPIISL305270" "EPIISL305271" "EPIISL305275" "EPIISL305277" "EPIISL305912" "EPIISL309495" "EPIISL309496" "EPIISL309497" "EPIISL309498" "EPIISL312940" "EPIISL312953" "EPIISL312956" "EPIISL313594" "EPIISL313595" "EPIISL313596" "EPIISL313597" "EPIISL316454" "EPIISL319781" "EPIISL319782" "EPIISL319783" "EPIISL319784" "EPIISL319785" "EPIISL319786" "EPIISL319787" "EPIISL319788" "EPIISL319789" "EPIISL319792" "EPIISL319793" "EPIISL319794" "EPIISL319795" "EPIISL319796" "EPIISL319799" "EPIISL319800" "EPIISL319801" "EPIISL319802" "EPIISL319803" "EPIISL319828" "EPIISL319829" "EPIISL319830" "EPIISL319831" "EPIISL319832" "EPIISL319833" "EPIISL319834" "EPIISL319835" "EPIISL319836" "EPIISL319837" "EPIISL319839" "EPIISL320256" "EPIISL320257" "EPIISL320258" "EPIISL320259" "EPIISL320260" "EPIISL320693" "EPIISL320710" "EPIISL320713" "EPIISL321969" "EPIISL322857" "EPIISL322858" "EPIISL322860" "EPIISL322861" "EPIISL322862" "EPIISL322863" "EPIISL322867" "EPIISL330360" "EPIISL330361" "EPIISL330362" "EPIISL330364" "EPIISL330366" "EPIISL330367" "EPIISL330368" "EPIISL330370" "EPIISL330371" "EPIISL330372" "EPIISL330374" "EPIISL330375" "EPIISL330377" "EPIISL330380" "EPIISL330381" "EPIISL330382" "EPIISL330383" "EPIISL330384" "EPIISL330385" "EPIISL330386" "EPIISL330388" "EPIISL330389" "EPIISL330390" "EPIISL330391" "EPIISL330392" "EPIISL330393" "EPIISL330394" "EPIISL330397" "EPIISL330823" "EPIISL330824" "EPIISL330825" "EPIISL330830" "EPIISL330836" "EPIISL330842" "EPIISL330843" "EPIISL330844" "EPIISL330847" "EPIISL330850" "EPIISL330851" "EPIISL330852" "EPIISL330853" "EPIISL330854" "EPIISL330856" "EPIISL330857" "EPIISL330858" "EPIISL330861" "EPIISL330864" "EPIISL331563" "EPIISL331564" "EPIISL331565" "EPIISL331566" "EPIISL331946" "EPIISL331951" "EPIISL331952" "EPIISL331956" "EPIISL332315" "EPIISL332397" "EPIISL332398" "EPIISL33576" "EPIISL335852" "EPIISL335854" "EPIISL335855" "EPIISL335858" "EPIISL335878" "EPIISL335879" "EPIISL335880" "EPIISL335883" "EPIISL335884" "EPIISL335911" "EPIISL335951" "EPIISL335952" "EPIISL335954" "EPIISL335955" "EPIISL336635" "EPIISL336636" "EPIISL336637" "EPIISL336638" "EPIISL336639" "EPIISL338706" "EPIISL34539" "EPIISL349812" "EPIISL349814" "EPIISL35013" "EPIISL35021" "EPIISL35022" "EPIISL351839" "EPIISL351842" "EPIISL351843" "EPIISL351848" "EPIISL351849" "EPIISL351874" "EPIISL353332" "EPIISL353346" "EPIISL353439" "EPIISL355491" "EPIISL355508" "EPIISL355524" "EPIISL355529" "EPIISL362255" "EPIISL362257" "EPIISL362284" "EPIISL364140" "EPIISL364141" "EPIISL364156" "EPIISL364554" "EPIISL377817" "EPIISL377818" "EPIISL390404" "EPIISL409210" "EPIISL409510" "EPIISL409511" "EPIISL409717" "EPIISL409736" "EPIISL409764" "EPIISL62231" "EPIISL62338" "EPIISL71367" "EPIISL78853" "EPIISL78854" "EPIISL78855" | "EPIISL130413" "EPIISL149681" "EPIISL171352" "EPIISL171359" "EPIISL171375" "EPIISL171376" "EPIISL171378" "EPIISL171380" "EPIISL171381" "EPIISL171394" "EPIISL171399" "EPIISL172531" "EPIISL172532" "EPIISL172592" "EPIISL172737" "EPIISL172750" "EPIISL172752" "EPIISL172755" "EPIISL172759" "EPIISL172773" "EPIISL172774" "EPIISL172776" "EPIISL172777" "EPIISL172779" "EPIISL172781" "EPIISL172784" "EPIISL172785" "EPIISL172787" "EPIISL172788" "EPIISL172799" "EPIISL172800" "EPIISL172802" "EPIISL172803" "EPIISL172808" "EPIISL172811" "EPIISL172814" "EPIISL174121" "EPIISL174127" "EPIISL174137" "EPIISL175197" "EPIISL175202" "EPIISL175210" "EPIISL175211" "EPIISL175217" "EPIISL176513" "EPIISL176515" "EPIISL176547" "EPIISL176548" "EPIISL176570" "EPIISL177536" "EPIISL177540" "EPIISL188871" "EPIISL189842" "EPIISL191672" "EPIISL191693" "EPIISL191694" "EPIISL192157" "EPIISL192158" "EPIISL192190" "EPIISL202451" "EPIISL203567" "EPIISL203587" "EPIISL203588" "EPIISL205282" "EPIISL205283" "EPIISL205284" "EPIISL206111" "EPIISL206166" "EPIISL206167" "EPIISL206171" "EPIISL206172" "EPIISL206173" "EPIISL206175" "EPIISL206177" "EPIISL206178" "EPIISL206179" "EPIISL206180" "EPIISL206181" "EPIISL206183" "EPIISL206184" "EPIISL206190" "EPIISL206191" "EPIISL206193" "EPIISL206194" "EPIISL206195" "EPIISL206196" "EPIISL206197" "EPIISL206199" "EPIISL206203" "EPIISL206204" "EPIISL206206" "EPIISL206207" "EPIISL206208" "EPIISL206211" "EPIISL206213" "EPIISL206214" "EPIISL206244" "EPIISL207025" "EPIISL207394" "EPIISL207395" "EPIISL207422" "EPIISL207431" "EPIISL207434" "EPIISL207455" "EPIISL207457" "EPIISL211679" "EPIISL211681" "EPIISL211682" "EPIISL211683" "EPIISL211685" "EPIISL211686" "EPIISL211687" "EPIISL211691" "EPIISL211693" "EPIISL211694" "EPIISL211705" "EPIISL211908" "EPIISL211910" "EPIISL211912" "EPIISL211919" "EPIISL211931" "EPIISL212061" "EPIISL212366" "EPIISL212370" "EPIISL212371" "EPIISL212999" "EPIISL213001" "EPIISL213003" "EPIISL213004" "EPIISL213005" "EPIISL213019" "EPIISL213975" "EPIISL213976" "EPIISL213977" "EPIISL213978" "EPIISL213979" "EPIISL215614" "EPIISL215615" "EPIISL215616" "EPIISL215617" "EPIISL215618" "EPIISL215619" "EPIISL215628" "EPIISL215629" "EPIISL215631" "EPIISL215632" "EPIISL215633" "EPIISL215635" "EPIISL215636" "EPIISL215770" "EPIISL215779" "EPIISL220268" "EPIISL220272" "EPIISL220275" "EPIISL220283" "EPIISL224270" "EPIISL225057" "EPIISL225058" "EPIISL225065" "EPIISL226798" "EPIISL230295" "EPIISL230296" "EPIISL230297" "EPIISL230298" "EPIISL230299" "EPIISL230300" "EPIISL230301" "EPIISL230302" "EPIISL230303" "EPIISL230304" "EPIISL230305" "EPIISL230306" "EPIISL230307" "EPIISL230308" "EPIISL230364" "EPIISL230365" "EPIISL230366" "EPIISL230368" "EPIISL230369" "EPIISL230372" "EPIISL230373" "EPIISL230375" "EPIISL232079" "EPIISL232093" "EPIISL232094" "EPIISL232096" "EPIISL232098" "EPIISL232540" "EPIISL232541" "EPIISL232543" "EPIISL232546" "EPIISL232555" "EPIISL232562" "EPIISL232564" "EPIISL233001" "EPIISL233726" "EPIISL235073" "EPIISL235075" "EPIISL235077" "EPIISL235088" "EPIISL235090" "EPIISL235091" "EPIISL235092" "EPIISL235093" "EPIISL235101" "EPIISL235104" "EPIISL235363" "EPIISL235537" "EPIISL249955" "EPIISL255341" "EPIISL255342" "EPIISL268306" "EPIISL268307" "EPIISL268308" "EPIISL268309" "EPIISL268310" "EPIISL268311" "EPIISL268313" "EPIISL268315" "EPIISL268316" "EPIISL268317" "EPIISL268318" "EPIISL268319" "EPIISL268320" "EPIISL268321" "EPIISL268322" "EPIISL268323" "EPIISL268324" "EPIISL268325" "EPIISL268326" "EPIISL268328" "EPIISL268331" "EPIISL270211" "EPIISL270212" "EPIISL273663" "EPIISL273665" "EPIISL273667" "EPIISL273671" "EPIISL273684" "EPIISL273686" "EPIISL275712" "EPIISL275928" "EPIISL275930" "EPIISL275932" "EPIISL275934" "EPIISL275936" "EPIISL275940" "EPIISL275943" "EPIISL275946" "EPIISL277139" "EPIISL278059" "EPIISL279044" "EPIISL279209" "EPIISL279214" "EPIISL279218" "EPIISL279252" "EPIISL279253" "EPIISL279254" "EPIISL281573" "EPIISL281685" "EPIISL281746" "EPIISL281747" "EPIISL281748" "EPIISL281749" "EPIISL281753" "EPIISL281755" "EPIISL281756" "EPIISL281757" "EPIISL281760" "EPIISL281771" "EPIISL281782" "EPIISL281802" "EPIISL281808" "EPIISL281817" "EPIISL281837" "EPIISL281848" "EPIISL281871" "EPIISL281877" "EPIISL281878" "EPIISL281882" "EPIISL281885" "EPIISL281888" "EPIISL282861" "EPIISL283312" "EPIISL285892" "EPIISL286127" "EPIISL286128" "EPIISL287619" "EPIISL288511" "EPIISL288512" "EPIISL288513" "EPIISL288515" "EPIISL288516" "EPIISL288517" "EPIISL288518" "EPIISL288519" "EPIISL290658" "EPIISL290659" "EPIISL290660" "EPIISL290661" "EPIISL292535" "EPIISL292537" "EPIISL292538" "EPIISL292539" "EPIISL296085" "EPIISL296523" "EPIISL299831" "EPIISL299834" "EPIISL299840" "EPIISL299842" "EPIISL299843" "EPIISL299852" "EPIISL299853" "EPIISL300840" "EPIISL304989" "EPIISL304990" "EPIISL304993" "EPIISL304994" "EPIISL304995" "EPIISL304996" "EPIISL306260" "EPIISL313517" "EPIISL319709" "EPIISL319711" "EPIISL320281" "EPIISL320344" "EPIISL320345" "EPIISL320346" "EPIISL320347" "EPIISL320348" "EPIISL320349" "EPIISL320351" "EPIISL320356" "EPIISL320360" "EPIISL322930" "EPIISL322934" "EPIISL322946" "EPIISL329853" "EPIISL330475" "EPIISL331067" "EPIISL331507" "EPIISL331761" "EPIISL331764" "EPIISL331766" "EPIISL331767" "EPIISL335644" "EPIISL335645" "EPIISL335646" "EPIISL335647" "EPIISL335648" "EPIISL335649" "EPIISL335656" "EPIISL336501" "EPIISL336503" "EPIISL336504" "EPIISL338561" "EPIISL338566" "EPIISL338619" "EPIISL338621" "EPIISL338622" "EPIISL338623" "EPIISL342243" "EPIISL346055" "EPIISL346056" "EPIISL346057" "EPIISL346059" "EPIISL346220" "EPIISL346222" "EPIISL347934" "EPIISL347940" "EPIISL347941" "EPIISL347944" "EPIISL347947" "EPIISL347950" "EPIISL347951" "EPIISL347952" "EPIISL348141" "EPIISL348142" "EPIISL348146" "EPIISL348160" "EPIISL350071" "EPIISL351879" "EPIISL352057" "EPIISL352058" "EPIISL353449" "EPIISL353513" "EPIISL353514" "EPIISL353543" "EPIISL355535" "EPIISL355536" "EPIISL355540" "EPIISL355541" "EPIISL355602" "EPIISL355604" "EPIISL355607" "EPIISL355634" "EPIISL355637" "EPIISL355762" "EPIISL356223" "EPIISL356224" "EPIISL356225" "EPIISL356292" "EPIISL356293" "EPIISL356294" "EPIISL356295" "EPIISL356296" "EPIISL356297" "EPIISL356299" "EPIISL356300" "EPIISL356303" "EPIISL356309" "EPIISL356311" "EPIISL356312" "EPIISL362170" "EPIISL362172" "EPIISL362173" "EPIISL362175" "EPIISL362177" "EPIISL362178" "EPIISL362179" "EPIISL362180" "EPIISL362181" "EPIISL362182" "EPIISL362188" "EPIISL362213" "EPIISL362214" "EPIISL362217" "EPIISL362220" "EPIISL362221" "EPIISL362222" "EPIISL362224" "EPIISL362245" "EPIISL362246" "EPIISL362250" "EPIISL362252" "EPIISL362254" "EPIISL362341" "EPIISL362389" "EPIISL362390" "EPIISL362391" "EPIISL362393" "EPIISL362394" "EPIISL362396" "EPIISL363855" "EPIISL363867" "EPIISL363869" "EPIISL363870" "EPIISL363871" "EPIISL363873" "EPIISL363875" "EPIISL363876" "EPIISL363877" "EPIISL363878" "EPIISL363880" "EPIISL363881" "EPIISL363884" "EPIISL363888" "EPIISL363892" "EPIISL363893" "EPIISL363895" "EPIISL363896" "EPIISL363897" "EPIISL363898" "EPIISL363899" "EPIISL363900" "EPIISL363901" "EPIISL363902" "EPIISL363904" "EPIISL363905" "EPIISL363906" "EPIISL363907" "EPIISL363957" "EPIISL363958" "EPIISL363962" "EPIISL363963" "EPIISL363968" "EPIISL363969" "EPIISL363970" "EPIISL363975" "EPIISL363976" "EPIISL364000" "EPIISL364001" "EPIISL364002" "EPIISL364003" "EPIISL364005" "EPIISL364010" "EPIISL364011" "EPIISL364540" "EPIISL364541" "EPIISL365767" "EPIISL365768" "EPIISL365769" "EPIISL365770" "EPIISL365771" "EPIISL365772" "EPIISL365773" "EPIISL365774" "EPIISL365775" "EPIISL365776" "EPIISL365778" "EPIISL365780" "EPIISL365781" "EPIISL365782" "EPIISL365783" "EPIISL365784" "EPIISL365785" "EPIISL365790" "EPIISL365834" "EPIISL365837" "EPIISL365974" "EPIISL366006" "EPIISL368194" "EPIISL368204" "EPIISL368206" "EPIISL377899" "EPIISL377900" "EPIISL377901" "EPIISL377902" "EPIISL377904" "EPIISL381578" "EPIISL381580" "EPIISL381582" "EPIISL381583" "EPIISL381584" "EPIISL381586" "EPIISL381588" "EPIISL381590" "EPIISL381592" "EPIISL381593" "EPIISL381598" "EPIISL381599" "EPIISL390122" "EPIISL390125" "EPIISL392520" "EPIISL392521" "EPIISL392523" "EPIISL392524" "EPIISL392526" "EPIISL393567" "EPIISL393571" "EPIISL393572" "EPIISL393574" "EPIISL393575" "EPIISL393576" "EPIISL393579" "EPIISL393581" "EPIISL393582" "EPIISL393583" "EPIISL393584" "EPIISL393588" "EPIISL393595" "EPIISL393596" "EPIISL393682" "EPIISL393684" "EPIISL393685" "EPIISL393686" "EPIISL393687" "EPIISL393688" "EPIISL393689" "EPIISL393690" "EPIISL393691" "EPIISL393692" "EPIISL393693" "EPIISL393694" "EPIISL393695" "EPIISL393696" "EPIISL393697" "EPIISL393698" "EPIISL393699" "EPIISL393700" "EPIISL393701" "EPIISL393702" "EPIISL393703" "EPIISL393705" "EPIISL393706" "EPIISL393707" "EPIISL393708" "EPIISL393709" "EPIISL393711" "EPIISL393712" "EPIISL393713" "EPIISL393714" "EPIISL393715" "EPIISL393717" "EPIISL393718" "EPIISL393719" "EPIISL393721" "EPIISL393723" "EPIISL393725" "EPIISL393726" "EPIISL393729" "EPIISL393730" "EPIISL393732" "EPIISL393733" "EPIISL393734" "EPIISL393736" "EPIISL393737" "EPIISL393738" "EPIISL393740" "EPIISL393742" "EPIISL393745" "EPIISL393748" "EPIISL393751" "EPIISL393757" "EPIISL393758" "EPIISL393759" "EPIISL393781" "EPIISL393782" "EPIISL393937" "EPIISL393938" "EPIISL393940" "EPIISL393941" "EPIISL393946" "EPIISL393955" "EPIISL393956" "EPIISL393960" "EPIISL393963" "EPIISL393967" "EPIISL394051" "EPIISL394052" "EPIISL394106" "EPIISL394107" "EPIISL394108" "EPIISL394110" "EPIISL394111" "EPIISL394112" "EPIISL394923" "EPIISL394924" "EPIISL394948" "EPIISL394949" "EPIISL394950" "EPIISL397182" "EPIISL397199" "EPIISL398311" "EPIISL398313" "EPIISL398314" "EPIISL398315" "EPIISL398532" "EPIISL398534" "EPIISL398535" "EPIISL398555" "EPIISL398790" "EPIISL398795" "EPIISL400779" "EPIISL400781" "EPIISL400782" "EPIISL400789" "EPIISL400792" "EPIISL400887" "EPIISL402407" "EPIISL402502" "EPIISL402504" "EPIISL402506" "EPIISL402508" "EPIISL402765" "EPIISL402766" "EPIISL402768" "EPIISL402769" "EPIISL402771" "EPIISL402772" "EPIISL402773" "EPIISL402774" "EPIISL402776" "EPIISL402779" "EPIISL402780" "EPIISL409343" "EPIISL409346" "EPIISL409350" "EPIISL409353" "EPIISL409356" "EPIISL409357" "EPIISL409358" "EPIISL409359" "EPIISL409361" "EPIISL409407" "EPIISL409410" "EPIISL409411" "EPIISL409412" "EPIISL409413" "EPIISL409414" "EPIISL409437" "EPIISL77944" "EPIISL84053" "EPIISL89055" "EPIISL89056" "EPIISL89058" "EPIISL89059" "EPIISL89060" "EPIISL89061" "EPIISL89063" "EPIISL89064" "EPIISL89065" "EPIISL89068" "EPIISL89069" "EPIISL89074" "EPIISL89077" "EPIISL89078" "EPIISL89079" "EPIISL89081" "EPIISL89082" "EPIISL89084" "EPIISL89085" "EPIISL89086" "EPIISL89090" "EPIISL89091" "EPIISL89092" "EPIISL89093" "EPIISL89097" "EPIISL89098" "EPIISL89100" "EPIISL89103" "EPIISL89105" "EPIISL89107" "EPIISL89111" "EPIISL89112" "EPIISL93638" |

**sTable 7:** **Accession numbers for influenza vaccine and clade reference strains downloaded from GISAID database (accessed on 27^th^ February 2020) and analysed in this study.**
